# Persistent telomere cohesion protects aged cells from premature senescence

**DOI:** 10.1101/2020.02.03.932145

**Authors:** Kameron Azarm, Amit Bhardwaj, Eugenie Kim, Susan Smith

## Abstract

Human telomeres are bound by the telomere repeat binding proteins TRF1 and TRF2. Telomere shortening in human cells leads to a DNA damage response that signals replicative senescence. While insufficient loading of TRF2 at shortened telomeres contributes to the DNA damage response in senescence, the contribution of TRF1 to senescence induction has not been determined. Here we show that counter to TRF2 deficiency-mediated induction of DNA damage, TRF1 deficiency serves a protective role to limit induction of DNA damage induced by subtelomere recombination. Shortened telomeres recruit insufficient TRF1 and as a consequence inadequate tankyrase 1 to resolve sister telomere cohesion. The persistent cohesion protects short telomeres from inappropriate recombination. Ultimately, in the final division, telomeres are no longer able to maintain cohesion and subtelomere copying ensues. Thus, the gradual loss of TRF1 and concomitant persistent cohesion that occurs with telomere shortening ensures a measured approach to replicative senescence.

## Introduction

Human telomeres contain long double-stranded arrays of TTAGGG repeats that end in a 3’ single-stranded overhang and are bound by the six-subunit shelterin complex ^1^. Due to the inability of DNA polymerases to replicate the ends of linear molecules and to nucleolytic processing that generates the 3’ overhang, telomeres shorten following each round of replication ^2, 3^. This shortening can be counteracted by telomerase, a reverse transcriptase which uses an RNA to add telomere repeats to the 3’ ends of chromosomes ^4, 5^. During human development telomerase is down regulated ^6^. As a result, human somatic cells undergo telomere shortening, which acts as a molecular clock inducing cells to cease division and senesce ^7^. The loss of telomeric DNA leads to insufficient chromosome-end protection and to activation of a DNA damage response (DDR) at telomeres that signals p53-dependent irreversible cell cycle arrest ^8^. The demonstration that introduction of telomerase can rescue the senescence phenotype indicates that it is due to telomere shortening ^9^.

Telomere function is regulated by the TTAGGG doubled-stranded repeat binding shelterin subunits TRF1 and TRF2. TRF1 plays a role in telomere replication by facilitating replication through the telomere repeats ^10^ and by regulating telomere length maintenance by telomerase ^11^. TRF1 function is aided by accessory binding proteins such as the poly(ADP-ribose) polymerase tankyrase 1 ^12, 13^, which in addition to a role in telomere length regulation, is required for resolution of sister telomere cohesion ^14, 15^. Cohesion between sister chromatids provides a template for recombination and repair during and after DNA replication. Maintenance of telomere cohesion in S phase relies on a third shelterin subunit TIN2 that binds to both TRF1 and TRF2 ^16^. Resolution of telomere cohesion relies on TRF1-mediated recruitment of tankyrase 1 to telomeres in G2 phase ^17, 18^.

TRF2 plays a distinct role at telomeres in chromosome end protection by facilitating formation of protective structures termed t-loops ^19–21^. Inactivation of TRF2 leads to a DDR at telomeres similar to that which occurs upon replicative senescence ^22^. The observation that overexpression of TRF2 can delay senescence onset suggests that the DDR in senescence is due to insufficient loading of shelterin at critically short telomeres ^23^. Loss of telomere function due to telomere shortening at a few chromosome ends is sufficient to induce arrest ^8, 24^. Quantitative analysis indicates that an aggregate of at least five DDR-positive telomeres is necessary to induce senescence ^25^. It has been suggested that telomeres fluctuate through different protective states depending on the level of TRF2; an intermediate state, which activates the DDR, but does not promote telomere fusion, would permit cells to cycle until they reach a threshold that activates a p53-dependent senescence arrest ^26, 27^.

Another feature of presenescent telomeres is that their resolution following DNA replication is delayed beyond S and G2 phase, into mitosis. Using fluorescent in situ hybridization (FISH) it was found that human telomeres remained unresolved (appeared as singlets) in metaphase in presenescent cells aged in culture and from early passage cells from aged individuals, whereas other chromosomal regions were resolved (appeared as doublets) ^28, 29^. The demonstration that introduction of telomerase into presenescent cells can rescue the metaphase telomere singlets suggests that they are due to telomere shortening ^29^. Depletion or knockout of tankyrase 1 leads to persistent telomere cohesion (FISH singlets in mitosis) ^14, 15^, reminiscent of the metaphase singlets in presenescent fibroblasts. However, the role of TRF1 and tankyrase 1 and the function of persistent cohesion in aged cells has not been determined.

Persistent telomere cohesion in mitosis has also been observed in ALT cells, telomerase negative cancer cells that maintain their telomeres through recombination-based mechanisms ^30^. In ALT cells persistent telomere cohesion was found to promote recombination between sister telomeres, while it suppressed inappropriate recombination between non-sisters ^31^. Although ALT cells have high levels of tankyrase 1 and TRF1, due to loss of ATRX (a common feature of ALT cells) ^32^, tankyrase 1 is sequestered away from telomeres ^31^. Overexpression of tankyrase 1 in ALT cells forces resolution of sister telomere cohesion and induces excessive subtelomere recombination between non-homologs, indicating a protective function for persistent telomere cohesion in ALT cancer cells ^31^. Whether persistent telomere cohesion plays a similar, protective role in senescing cells has not been determined.

Here we show that insufficient loading of TRF1 at shortened telomeres protects aged cells from an abrupt recombination-induced DDR. We demonstrate that it is the shortened telomeres *per se* in any cellular context (aged normal cells, ALT cancer cells, or telomerase-inhibited telomerase positive cancer cells) that induce persistent telomere cohesion, providing an inherent protective mechanism that accompanies telomere shortening. Specifically, in aged cells, the gradual loss of telomere repeats concomitant with TRF1 deficiency-induced persistent cohesion, contributes to attenuated onset of senescence.

## Results

### TRF1-mediated recruitment of tankyrase controls resolution of cohesion and subtelomere recombination in aged human fibroblasts

Tankyrase 1 binds to TRF1 through a consensus tankyrase binding site RGCADG in the amino terminal acidic domain of TRF1 (Fig. S1A)^13, 33^. To determine if resolution of telomere cohesion depends specifically on the tankyrase binding site in TRF1, we used CRISPR/Cas9 to mutate the essential terminal G of the RGCAD<u>G</u> site in the endogenous TRF1 gene to P (TRF1.G18P) in HEK293T cells (Fig. S1A). Multiple independent clones were isolated and sequenced (Fig. S1B) and telomere cohesion analyzed by isolating mitotic cells using mechanical shake-off and probing with a chromosome specific subtelomere probe 16p (triploid in HEK293T cells). As shown in Fig. 1A and 1B, a WT clone displayed normal resolution of telomere cohesion (doublets), whereas the three TRF1.G18P mutant clones showed persistent telomere cohesion (singlets), confirming that resolution of telomere cohesion depends on the tankyrase binding site in TRF1.

**Figure 1.**
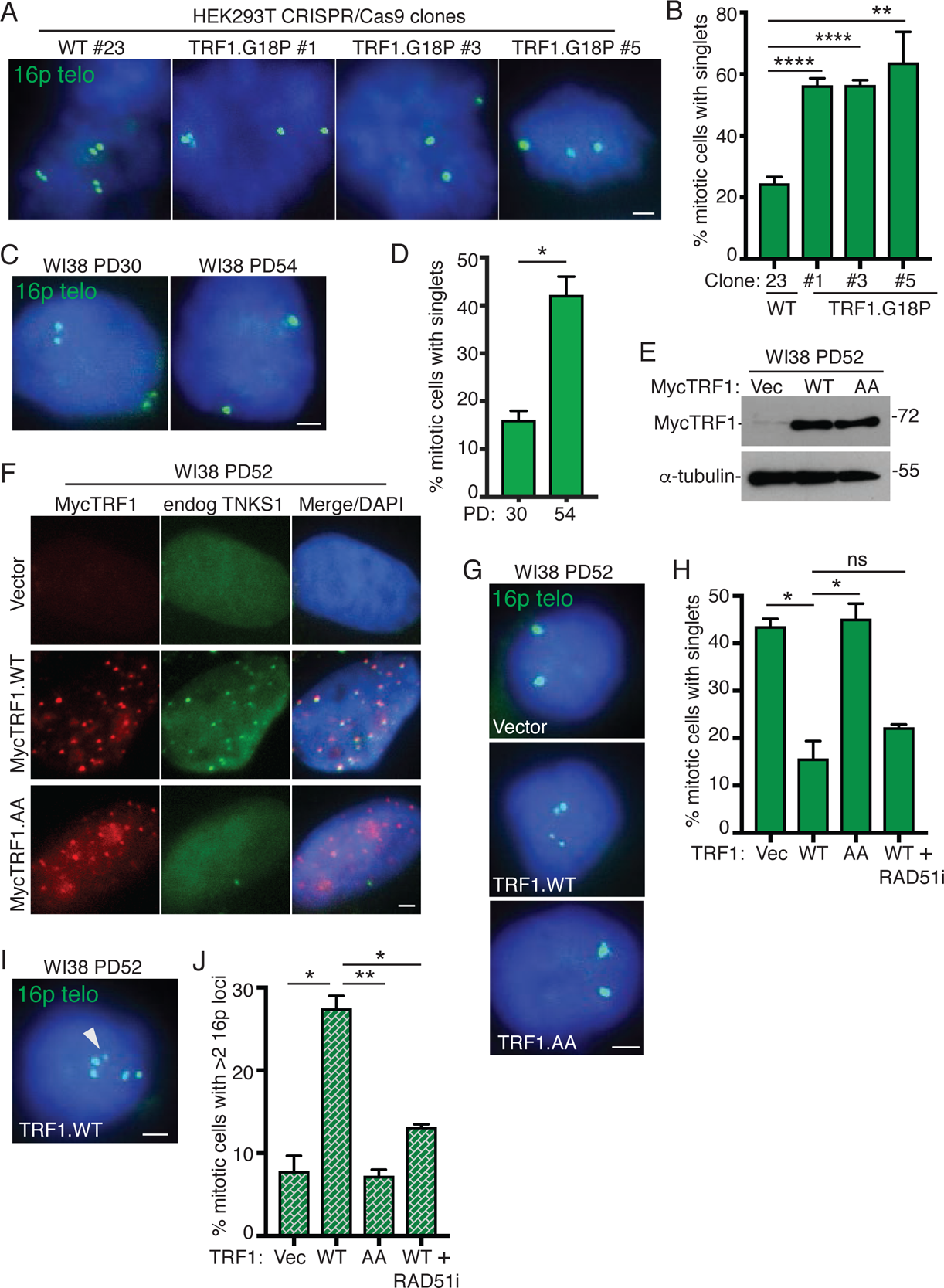
**TRF1-mediated recruitment of tankyrase 1 controls resolution of cohesion and suppresses Rad51-dependent subtelomere recombination in aged human fibroblasts.** **(A)** FISH analysis of HEK293T wild-type (#23) and TRF1.G18P mutant (#1, #3, #5) mitotic cells with a 16p (triploid in HEK293T cells) telo probe (green). **(B)** Quantification of the frequency of mitotic cells with cohered telomeres. Average of three independent experiments (n=28-69 cells each) ± SEM. **(C)** FISH analysis of early (PD30) and late (PD54) passage WI38 mitotic cells with a 16p telo probe (green). **(D)** Quantification of the frequency of mitotic cells with cohered telomeres. Average of two independent experiments (n=36-50 cells each) ± SEM. **(E)** Immunoblot analysis of Vector, TRF1.WT, or TRF1.AA transfected late (PD52) WI38 cell extracts. **(F)** Immunofluorescence analysis of Vector, TRF1.WT, or TRF1.AA transfected late (PD52) WI38 cells using Myc (red) and TNKS1 (green) antibodies. **(G)** FISH analysis of Vector, TRF1.WT, or TRF1.AA transfected late (PD52) WI38 mitotic cells using a 16p telo probe (green). **(H)** Quantification of the frequency of mitotic cells with cohered telomeres. Average of two independent experiments (n=33-50 cells each) ± SEM. **(I)** FISH analysis of a TRF1.WT transfected late (PD52) WI38 mitotic cell exhibiting subtelomere copying (arrowhead) using a 16p telo probe (green). **(J)** Quantification of the frequency of mitotic cells exhibiting subtelomere copying. Average of two independent experiments (n=33-50 cells each) ± SEM. (A, C, F, G, I) DNA was stained with DAPI (blue). Scale bars represent 2 μm. *p ≤ 0.05, **p ≤ 0.01, ****p ≤ 0.0001, Student’s unpaired t test. See also Figure S1. Source data are provided as a Source Data file.

As cells approach replicative senescence they exhibit persistent telomere cohesion, shown in Fig. 1C and 1D for aged WI38 cells and previously ^28, 29, 34^. During physiological telomere shortening shelterin components become limiting. Immunofluorescence analysis shows a decrease in TRF1 at aged cell telomeres (Fig. S1C). We thus asked if there was insufficient TRF1 on aged cell telomeres to recruit tankyrase 1 for resolution of telomere cohesion. Overexpression of wild type TRF1 (TRF1.WT) by transient transfection (20 hr) in aged WI38 cells (Fig. 1E) led to its accumulation on telomeres and to recruitment of endogenous tankyrase 1 to telomeres (Fig. 1F and S1D), whereas overexpression of a mutant allele, TRF1.AA, where the essential terminal G (and adjacent D) in the RGCAD<u>G</u> tankyrase binding site was mutated to A (Fig. S1E) ^18, 35^, similarly led to its accumulation on telomeres, but not to recruitment of endogenous tankyrase 1 (Fig. 1F and S1D). To determine if the recruitment of excess tankyrase 1 to telomeres was sufficient to force resolution of cohesion, we performed 16P subtelomere FISH analysis. As shown in Fig. 1G and 1H, TRF1.WT, but not Vector or TRF1.AA, forced resolution of cohesion in aged WI38 fibroblasts. Similar results were obtained in aged IMR90 cells (Fig. S1F-S1H). Finally, FISH analysis with a dual 13q subtelomere/arm probe showed similar results for the subtelomere (Fig. 1I).

Previous studies showed that forcing resolution of cohesion in ALT cancer cells led to RAD51-dependent subtelomere recombination between nonhomologous sisters evidenced by an increase in the number of 16p subtelomere loci ^31^. FISH analysis indicated a significant increase in the frequency of mitotic cells with greater than two 16p loci in aged WI38 cells transfected with TRF1.WT, but not Vector or TRF1.AA (Fig. 1I and 1J), indicating that forced resolution of cohesion leads to subtelomere recombination in aged cells. Similar results were obtained in aged IMR90 cells (Fig. S1J and S1K) and FISH analysis with the dual 13q subtelomere/arm probe showed that recombination was specific to the subtelomere (Fig. 1L). To determine if the observed subtelomere recombination was dependent on RAD51, TRF1.WT transfected cells were treated with a RAD51 small molecule inhibitor (RAD51i). Resolution of telomere cohesion was unaffected by inhibition of RAD51 (Fig. 1H), however subtelomere recombination was abrogated (Fig. 1J), indicating that forced resolution of cohesion by overexpression of TRF1 leads to RAD51-dependent subtelomere recombination in aged cells.

To ascertain additional requirements for subtelomere recombination, we forced resolution of cohesion with TRF1.WT and interrogated cells with multiple small molecule inhibitors and siRNAs (Fig. 2A– 2C). Resolution of cohesion occurred under all conditions (Fig. 2A) demonstrating that the treatments did not inhibit resolution. However, subtelomere copying was inhibited in cells treated with ATR or CHK1 inhibitors (Fig. 2B). The requirement for CHK1 and ATR, along with RAD51 (shown in Fig. 1J) suggests a homologous recombination mechanism for subtelomere copying. Recent studies in ALT cancer cells found that telomere recombination can proceed through multiple mechanisms, including POLD3-dependent break-induced telomere synthesis ^36–38^ and RAD52-dependent mitotic telomeric DNA synthesis ^39^. However, the observation that subtelomere copying does not depend on RAD52, CHK2, or POLD3 (Fig. 2B) indicates that it is distinct from these ALT telomere recombination pathways.

**Figure 2.**
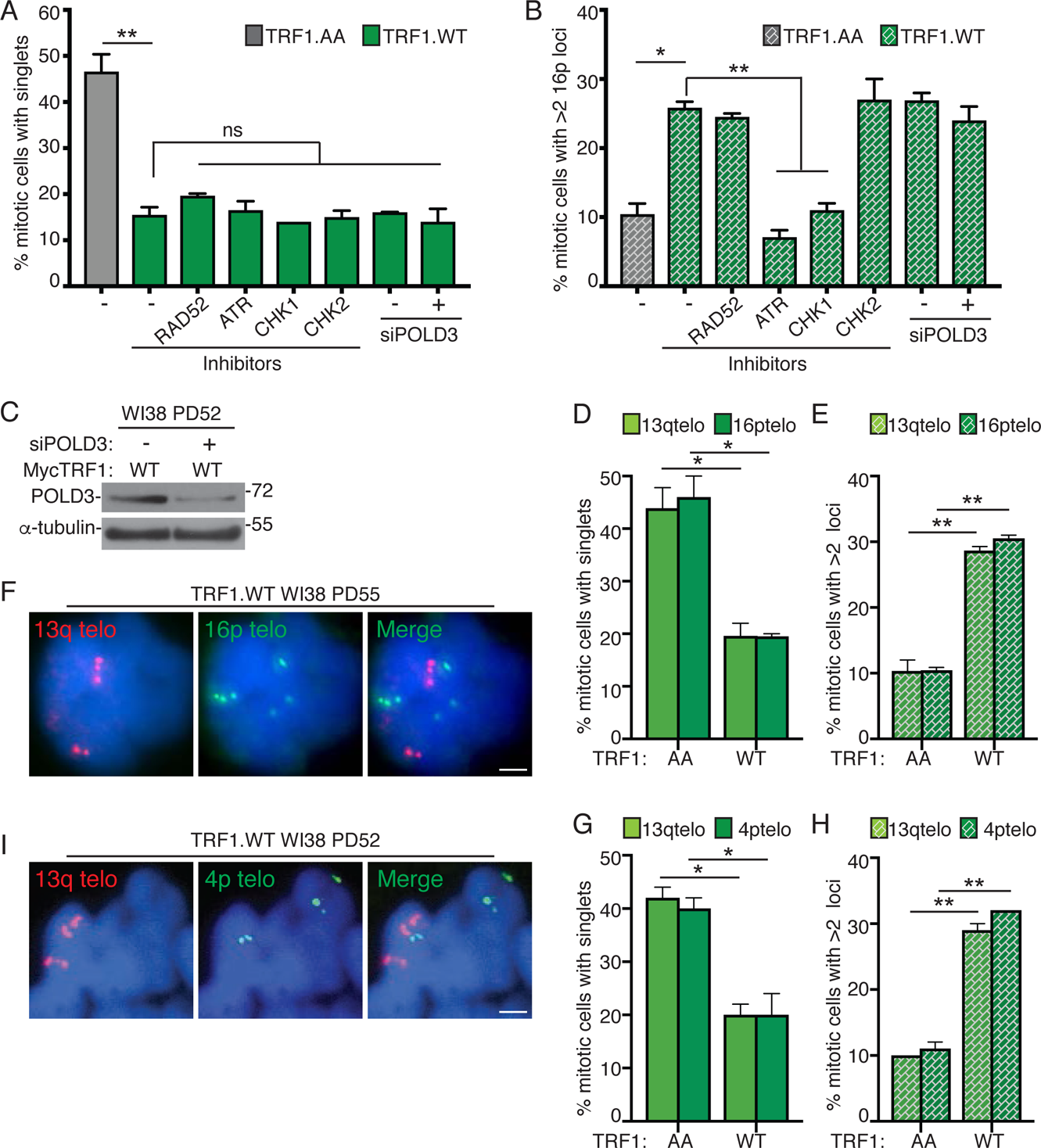
Rad51-dependent subtelomere recombination triggered by forced resolution of cohesion is ATR and ChkI-dependent, POLD3-independent, and involves multiple subtelomeres. (A and B) Quantification of the frequency of TRF1.AA or TRF1.WT transfected late (PD50-53) WI38 mitotic cells (A) with cohered telomeres or (B) exhibiting subtelomere copying measured by FISH analysis with a 16p telo probe following treatment with the indicated inhibitors or siRNA. For TRF1.AA transfection (n=46). For TRF1.WT transfection, average of 2 independent experiments (n=25-50 cells each) ± SEM. (C) Immunoblot analysis of TRF1.WT-transfected, POLD3 siRNA-treated late (PD52) WI38 cell extracts. (D and E) Quantification of the frequency of WI38 TRF1.AA or TRF1.WT transfected late (PD55) mitotic cells (D) with cohered telomeres or (E) exhibiting subtelomere copying measured by dual FISH analysis with 13q and 16p telo probes. Average of two independent experiments (n=46-58 cells each) ± SEM. (F) FISH analysis of a WI38 TRF1.WT transfected late (PD55) mitotic cell exhibiting subtelomere copying using 13q (red) and 16p (green) telo probes. (G and H) Quantification of the frequency of TRF1.AA or TRF1.WT transfected late (PD52) WI38 mitotic cells (G) with cohered telomeres or (H) exhibiting subtelomere copying measured by dual FISH analysis with 13q and 4p telo probes. Average of two independent experiments (n=50 cells each) ± SEM. (I) FISH analysis of a TRF1.WT transfected late (PD52) WI38 mitotic cell exhibiting subtelomere copying using 13q (red) and 4p (green) telo probes. (F and I) DNA was stained with DAPI (blue). Scale bars represent 2 μm. *p ≤ 0.05, **p ≤ 0.01, (ns) not significant, Student’s unpaired t test. Source data are provided as a Source Data file.

Finally, we asked if other subtelomeres (in addition to 16p) underwent copying by performing dual FISH analysis with subtelomere probes 13q and 16p (Fig. 2D-2F) or 13q and 4p (Fig. 2G–2I). Regardless of the probe used, aged WI38 cells transfected with TRF1.WT (but not TRF1.AA) showed a similar reduction in telomere cohesion (Fig. 2D and 2G) and a similar increase in subtelomere copying (Fig. 2E and 2H). To determine if multiple telomeres were undergoing recombination in the same cell, we asked what percentage of cells undergoing 13q copying also showed 16p or 4p copying. We found that 61% of TRF1.WT transfected cells that showed 13q copying also exhibited 16p copying (see Fig. 2F for example). Similarly, 72% of TRF1.WT transfected cells that showed 13q copying also showed 4p copying (see Fig. 2I for example). These data indicate that multiple subtelomeres undergo recombination in the same cell and (since the analysis is done within 20 hr of transfection) in the same cell cycle.

### Subtelomere recombination leads to DNA damage and premature senescence

As the RAD51-dependent subtelomere recombination appeared to occur at high frequency and at multiple subtelomeres, we asked if we could detect an increase in RAD51 foci upon forced resolution of cohesion. Overexpression of TRF1.WT, but not Vector or TRF1.AA, following lentiviral infection in aged WI38 cells (Fig. S2A) led to a greater than three-fold increase in RAD51 foci overall (Fig. 3A and 3B), and a greater than four-fold increase in RAD51 foci associated with telomeres (Fig. 3C and 3D). To determine if the observed high level of subtelomere recombination coincided with a DNA damage response, we use immunofluorescence analysis to measure the levels of γH2AX and 53BP1. Overexpression of TRF1.WT, but not Vector or TRF1.AA led to a two-fold increase in DNA damage foci (Fig. 3E and 3F), and a three-fold increase in γH2AX foci associated with telomeres (Fig. 3G and 3H). Treatment of TRF1.WT overexpressing cells with the RAD51 inhibitor abrogated the increase in DNA damage (Fig. 3F). As we showed above that RAD51 is required for subtelomere recombination, but not for resolution of cohesion, these results indicate that subtelomere recombination drives the DNA damage.

**Figure 3.**
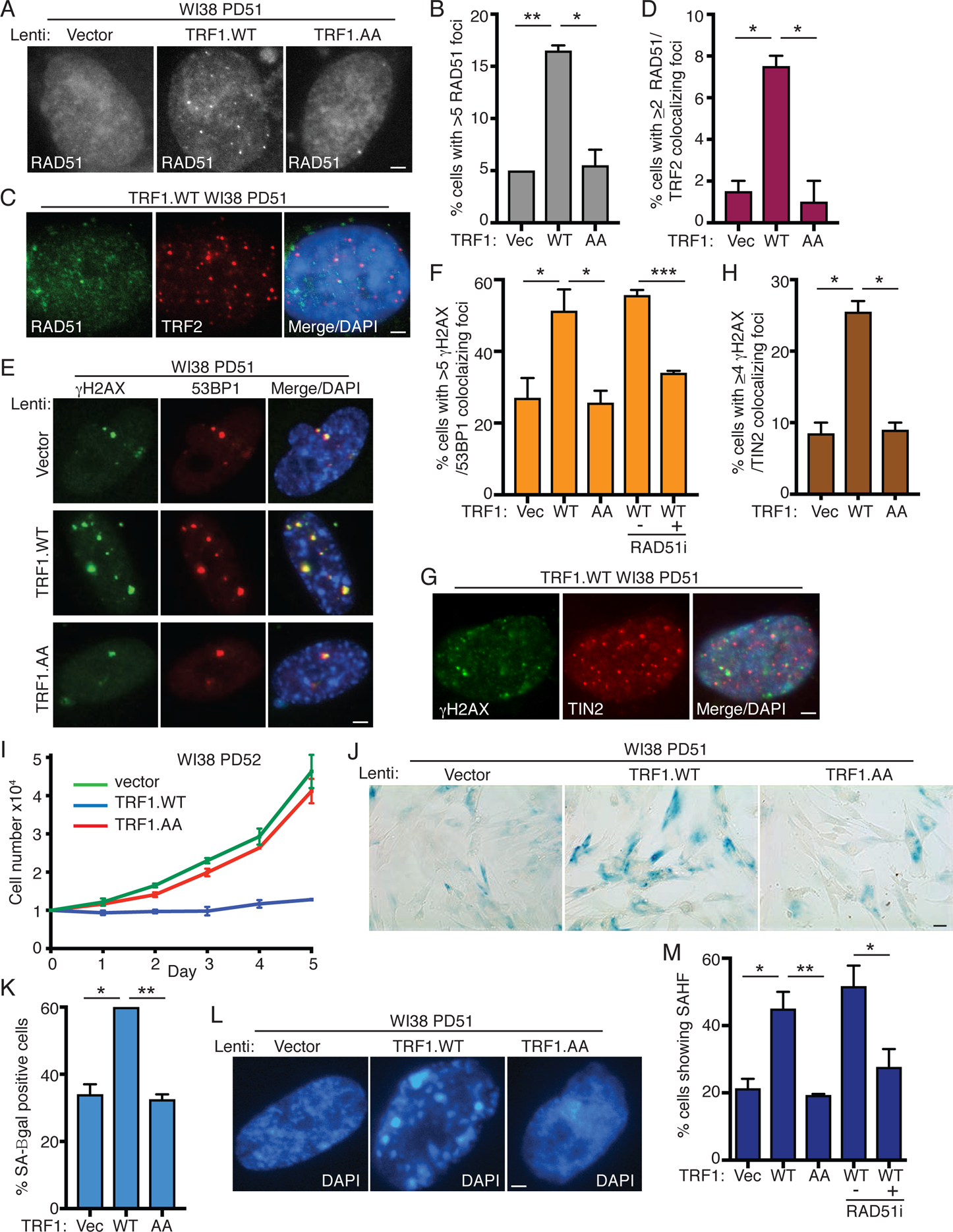
Subtelomere recombination leads to DNA damage and premature senescence in aged human fibroblasts. **(A)** Immunofluorescence analysis of Vector, TRF1.WT, or TRF1.AA infected late (PD51) WI38 cells with RAD51 antibody. **(B)** Quantification of the frequency of cells displaying >5 RAD51 foci. Average of two independent experiments (n=100 cells each) ± SEM. **(C)** Immunofluorescence analysis of TRF1.WT-infected late (PD51) WI38 cells with Rad51 (green) and TRF2 (red) antibodies. **(D)** Quantification of the frequency of cells displaying ≥2 Rad51/TRF2 colocalizations. Average of two independent experiments (n=100 cells each) ± SEM. **(E)** Immunofluorescence analysis of Vector, TRF1.WT, or TRF1.AA infected late (PD51) WI38 cells with γH2AX (green) and 53BP1 (red) antibodies. **(F)** Quantification of the frequency of cells displaying >5 γH2AX/53BP1 colocalizing foci. Average of three independent experiments (n=55-100 cells each) ± SEM. **(G)** Immunofluorescence analysis of TRF1.WT-infected late (PD51) WI38 cells with γH2AX (green) and TIN2 (red) antibodies. **(H)** Quantification of the frequency of cells with ≥4 γH2AX/TIN2 colocalizing foci. Average of two independent experiments (n=100 cells each) ± SEM. **(I)** Growth curve analysis of Vector, TRF1.WT, or TRF1.AA infected late (PD52) WI38 cells. Average of two independent experiments ± SEM. **(J)** SA-β-gal analysis of Vector, TRF1.WT, or TRF1.AA infected late (PD51) WI38 cells. Scale bar represent 100 μm. **(K)** Quantification of SA-β-gal positive cells. Average of two independent experiments (n=50-100 cells each) ± SEM. **(L)** Detection of senescence-associated heterochromatin foci (SAHF) in Vector, TRF1.WT, or TRF1.AA infected late (PD51) WI38. **(M)** Quantification of SAHF-positive cells. Average of three independent experiments (n=55-100 cells each) ± SEM. (C, E, G, and L) DNA was stained with DAPI (blue). (A, C, E, G, L) Scale bars represent 2 μm. *p ≤ 0.05, **p ≤ 0.01, ***p ≤ 0.001, Student’s unpaired t test. See also Figure S2. Source data are provided as a Source Data file.

To determine the impact of forced resolution of telomere cohesion on cell growth, we performed growth curve analysis following lentiviral infection of aged WI38 cells with TRF1 alleles. As shown in Fig. 3I, we observed a dramatic growth arrest specifically in TRF1.WT, but not Vector or TRF1.AA expressing cells. Introduction of the same TRF1 alleles into young WI38 cells had no effect on cell growth (Fig. S2B and S2C), as expected and consistent with previous studies ^23^. Inhibition of cell growth was due to premature activation of senescence as evidenced by an increase in the senescence associated marker β-galactosidase SA-β-gal ^40^ (Fig.3J and 3K) and in senescence associated heterochromatin foci (SAHF) ^41^ (Fig. 3L and 3M) specifically in TRF1.WT cells. The premature senescence was abrogated by treatment of TRF1.WT cells with the RAD51 inhibitor (Fig. 3M), consistent with a role for subtelomere recombination-induced DNA damage in induction of premature senescence.

### Loss of the DNA damage checkpoint rescues TRF1-induced premature senescence and leads to runaway subtelomere recombination

Next, to determine if induction of premature senescence was checkpoint dependent, we used SV40 Large T (LT) antigen to extend the proliferative life span of aged fibroblasts. We introduced a Vector or SV40-LT into WI38 cells at PD42 by retroviral infection. Once established, we allowed the cell lines to continue to grow until we observed slowing (pre senescence) of the Vector control cells, while SV40-LT cells continued to proliferate (PD6) (Fig. 4A). Telomere cohesion was measured at PD6 to determine if life-span extension interfered with persistent cohesion. As shown in Fig. 4B and 4C, both Vector and SV40-LT cells exhibited similar levels of persistent cohesion. However, SV40-LT cells showed an increase in subtelomere recombination compared to Vector (Fig. 4D), suggesting that loss of the checkpoint permits a low level of subtelomere copying even with persistent cohesion.

**Figure 4.**
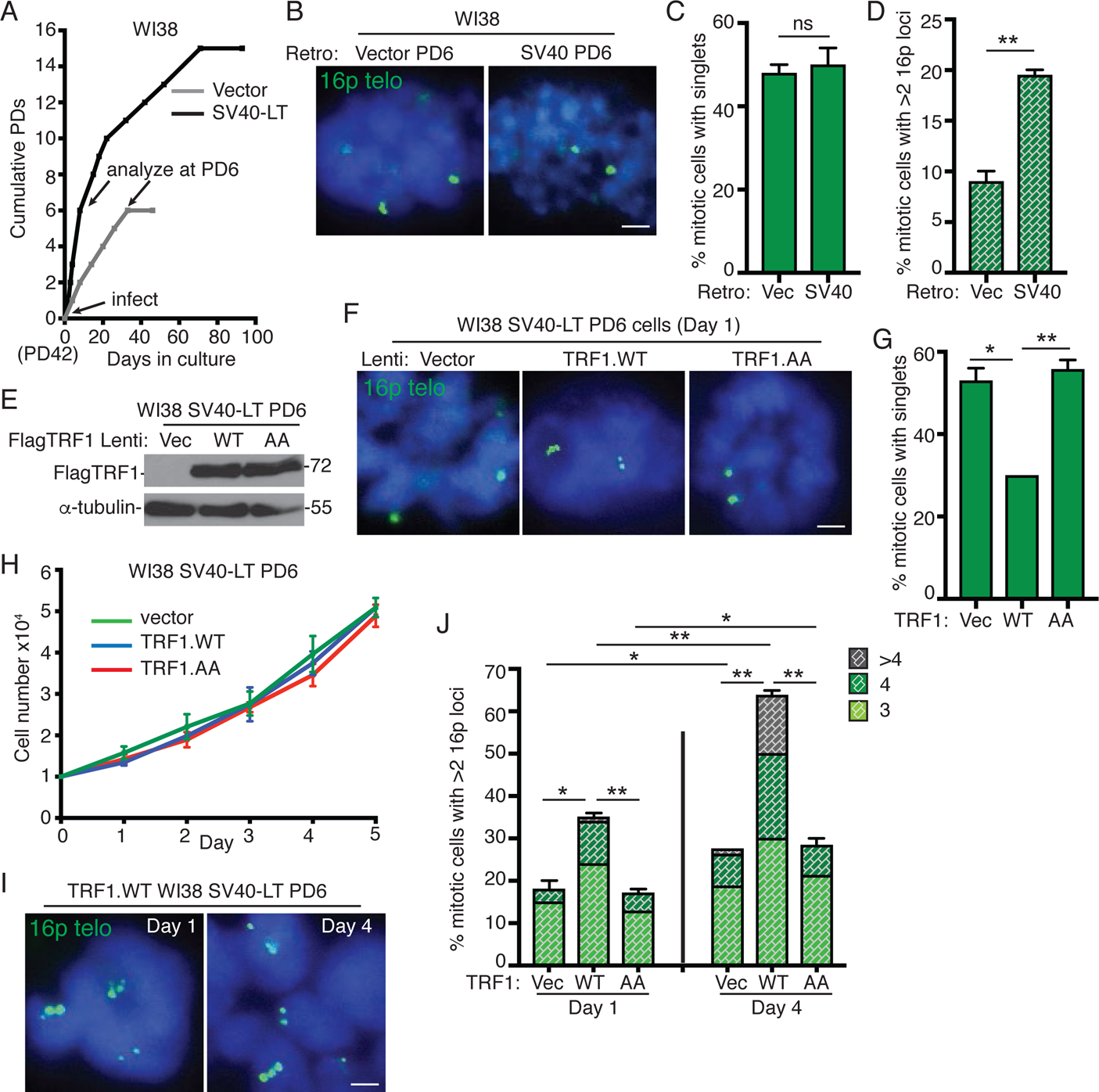
Loss of the DNA damage checkpoint rescues TRF1-induced premature senescence and leads to runaway subtelomere recombination. **(A)** Growth curve analysis of Vector or SV40-LT infected mid (PD42) WI38 cells. Subsequent analyses were performed at PD6 (indicated) when Vector-infected cells begin to senesce. **(B)** FISH analysis of Vector or SV40-LT infected WI38 mitotic cells at PD6 using a 16p telo probe (green). (C and D) Quantification of the frequency of mitotic cells (C) with cohered telomeres or (D) exhibiting subtelomere copying in Vector or SV40-LT-infected WI38 cells at PD6. Average of two independent experiments (n=41-60 cells each) ± SEM. **(E)** Immunoblot analysis of Vector, TRF1.WT, or TRF1.AA infected WI38 SV40-LT (PD6) cell extracts. **(F)** FISH analysis of Vector, TRF1.WT, or TRF1.AA infected WI38 SV40-LT (PD6) cells on Day 1 of growth curve analysis (H) using a 16p telo probe (green). **(G)** Quantification of the frequency of mitotic cells with cohered telomeres. Average of two independent experiments (n=43-50 cells each) ± SEM. **(H)** Growth curves of Vector, TRF1.WT, or TRF1.AA WI38 infected SV40-LT (PD6) cells. Average of two independent experiments ± SEM. **(I)** FISH analysis of TRF1.WT infected WI38 SV40-LT (PD6) mitotic cells exhibiting subtelomere copying on Day 1 and Day 4 of growth curve analysis (F) using a 16p telo probe (green). **(J)** Quantification of the frequency of mitotic cells exhibiting subtelomere copying. Average of two independent experiments (n=40-50 cells each) ± SEM. (B, G, I) DNA was stained with DAPI (blue). Scale bars represent 2 μm. *p ≤ 0.05, **p ≤ 0.01, Student’s unpaired t test. See also Figure S3. Source data are provided as a Source Data file.

We next introduced the Vector, TRF1.WT, and TRF1.AA alleles by lentiviral infection into WI38 SV40-LT PD6 cells (Fig. 4E). FISH analysis showed that TRF1.WT, but not Vector or TRF1.AA, forced resolution of telomere cohesion (Fig. 4F and 4G), as it had in aged WI38 cells. However, in contrast to aged WI38 cells, TRF1.WT-infected WI38 SV40-LT cells did not undergo growth arrest; they proliferated at the same rate as Vector or TRF1.AA cells (Fig. 4H), indicating that loss of the checkpoint abrogated the growth arrest. Since these cells continued to grow, we were able to analyze subtelomere copying at the standard early time point after infection (day 1) as well as later at day 4. TRF1.WT cells showed subtelomere copying on day 1 that increased dramatically (in frequency and number of subtelomeres copied) on day 4, indicating runaway telomere copying in the absence of checkpoint-mediated growth arrest (Fig. 4I and 4J). FACS analysis showed that the increase in copying was not due to an increase in ploidy (Fig. S3A). We also observed a slighter but significant increase in subtelomere copying in Vector and TRF1.AA cells from day 1 to day 4 (Fig. 4J), again indicating that loss of the checkpoint reveals a low level of subtelomere copying even with persistent cohesion.

### TRF1 overexpression in ALT cancer cells induces subtelomere recombination and activates senescence

Our studies thus far in normal primary cells suggest that telomere shortening that occurs in the absence of telomerase induces persistent telomere cohesion (resulting from the inability of limiting TRF1 to recruit tankyrase 1), which serves as a protective mechanism against subtelomere recombination, DNA damage, and premature activation of senescence. We sought to determine if a similar mechanism was at work in ALT cancer cells, which (like aged fibroblasts) exhibit persistent telomere cohesion. We previously identified an ATRX–macroH2A1.1– tankyrase axis, where the absence of ATRX freed the soluble macroH2A1.1 pool to sequester tankyrase 1 away from telomeres; we showed that overexpression of ATRX or tankyrase 1 in ALT cells forced resolution of telomere cohesion and led to Rad51-dependent subtelomere recombination ^31^. Although ALT cells have exceptionally long telomeres, they also harbor critically short ones due to the absence of telomerase ^42^. We thus asked if TRF1 was limiting for resolution of telomere cohesion in ALT cells. We introduced Vector, TRF1.WT, or TRF1.AA into GM847 ALT cells by transient transfection and performed immunoblot (Fig. 5A) and immunofluorescence (Fig. 5B) analysis.

**Figure 5.**
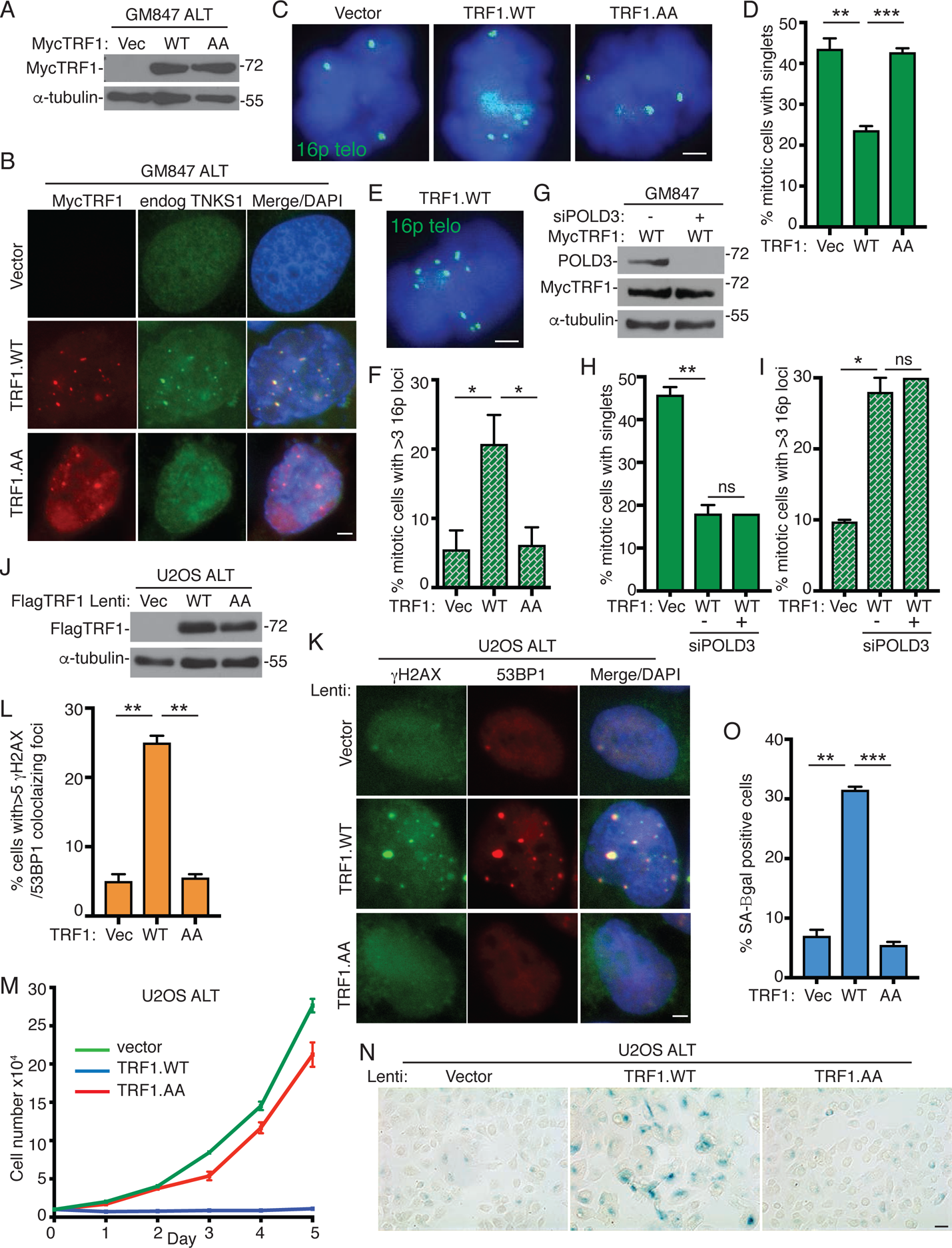
TRF1 overexpression in ALT cancer cells induces subtelomere recombination and activates senescence. **(A)** Immunoblot analysis of Vector, TRF1.WT, or TRF1.AA transfected GM847 cell extracts. **(B)** Immunofluorescence analysis of Vector, TRF1.WT, or TRF1.AA transfected GM847 cells using Myc (red) and TNKS1 (green) antibodies. **(C)** FISH analysis of Vector, TRF1.WT, or TRF1.AA transfected GM847 mitotic cells using a 16p (triploid in GM847 cells) telo probe (green). **(D)** Quantification of the frequency of mitotic cells with cohered telomeres. Average of three independent experiments (n=30-42 cells each) ± SEM. **(E)** FISH analysis of a TRF1.WT transfected GM847 mitotic cell exhibiting subtelomere copying using a 16p telo probe (green). **(F)** Quantification of the frequency of mitotic cells exhibiting subtelomere copying. Average of three independent experiments (n=40-42 cells each) ± SEM. **(G)** Immunoblot analysis of TRF1.WT-transfected, POLD3 siRNA-treated GM847 cell extracts. (H and I) Quantification of the frequency of mitotic cells (H) with cohered telomeres or (I) exhibiting subtelomere copying. Average of two independent experiments (n=50 cells each) ± SEM. **(J)** Immunoblot analysis of Vector, TRF1.WT, or TRF1.AA infected U2OS cell extracts. **(K)** Immunofluorescence analysis of Vector, TRF1.WT, or TRF1.AA infected U2OS cells with γH2AX (green) and 53BP1 (red) antibodies. **(L)** Quantification of the frequency of cells with >5 γH2AX/53BP1 colocalizing foci. Average of two independent experiments (n=100 cells each) ± SEM. **(M)** Growth curve analysis of Vector, TRF1.WT, or TRF1.AA infected U2OS cells. Average of three technical replicates ± SEM. **(N)** SA-β-gal analysis of Vector, TRF1.WT, or TRF1.AA infected U2OS cells. Scale bar represent 100 μm. **(O)** Quantification of SA-β-gal positive cells. Average of two independent experiments (n=50-77 cells each) ± SEM. (B, C, E, K) DNA was stained with DAPI (blue). Scale bars represent 2 μm. *p ≤ 0.05, **p ≤ 0.01, ***p ≤ 0.001, (ns) not significant, Student’s unpaired t test. See also Figure S4 Source data are provided as a Source Data file.

Both TRF1 alleles accumulated on ALT telomeres, but only TRF1.WT (not TRF1.AA) recruited endogenous tankyrase 1 (Fig. 5B and S4A). FISH analysis with the 16p subtelomere probe (triploid in GM847 cells) showed that overexpression of TRF1.WT, but not Vector or TRF1.AA, led to resolution of persistent cohesion (Fig. 5C and 5D) and subtelomere recombination (Fig. 5E and 5F). Analysis of metaphase spreads showed that upon forced resolution of cohesion subtelomere sequences can be detected on additional chromosomes, consistent with subtelomere copying (Fig. S4B-S4E). Subtelomere recombination was independent of POLD3 (Fig. 5G-5I), indicating that subtelomere copying in ALT cells (as in aged cells) is distinct from POLD3-dependent break-induced telomere synthesis.

We next asked if forced resolution of cohesion and subtelomere recombination would have the same consequences in ALT cells as in normal aged cells, where we showed above that subtelomere recombination led to DNA damage and checkpoint-dependent premature senescence. We used lentiviral infection to introduce the vector, TRF1.WT, and TRF1.AA alleles into U2OS, a checkpoint proficient ALT cell line with wild-type p53 (immunoblot in Fig. 5J). Immunofluorescence analysis showed that TRF1.WT (but not Vector or TRF1.AA) led to induction of DNA damage (Fig. 5K and 5L) and to a growth arrest (Fig. 5M) due to activation of senescence, indicated by an increase in SA-β-gal positive cells (Fig. 5N and 5O), similar to the effect of TRF1.WT in aged cells and consistent with a checkpoint-mediated growth arrest. Introduction of the same alleles into GM847, an ALT cell line lacking p53 checkpoint function, did not lead to a growth arrest (Fig. S4F).

### Telomerase overrides the need for persistent telomere cohesion and suppresses subtelomere recombination

Our data indicate that TRF1 is limiting for telomere resolution in two cell types that lack telomerase (ALT cancer cells and normal primary cells) suggesting that short telomeres drive the persistent cohesion. Previous studies showed that introduction of telomerase into normal aged cells rescued persistent telomere cohesion ^29^. We thus asked if telomerase could rescue persistent telomere cohesion in ALT cells. We introduced TERT/TR, a vector expressing the wild type (WT) telomerase catalytic subunit and RNA into GM847 ALT cells by transient transfection and subjected the cells to immunoblot (Fig. 6A) and FISH (Fig. 6B) analysis. Introduction of telomerase rescued persistent telomere cohesion in ALT cells (Fig. 6B and 6C). Rescue depended on the catalytic activity of telomerase; a catalytically dead (CD) TERT/TR did not rescue persistent telomere cohesion (Fig. 6B and 6C). Interestingly, while telomerase expression (like TRF1 overexpression) forced resolution of cohesion in ALT cells, (unlike TRF1 overexpression), it did not lead to subtelomere recombination (Fig. 6D). Similar results were obtained upon introduction of telomerase into ALT U2OS cells (Fig. S5A-S5D). The observation that telomerase can rescue persistent cohesion suggests that it is the critically short telomeres in ALT cells that give rise to persistent cohesion. The absence of subtelomere recombination upon forced resolution by telomerase further suggests that it is the critically short telomeres that are responsible for the subtelomere recombination.

**Figure 6.**
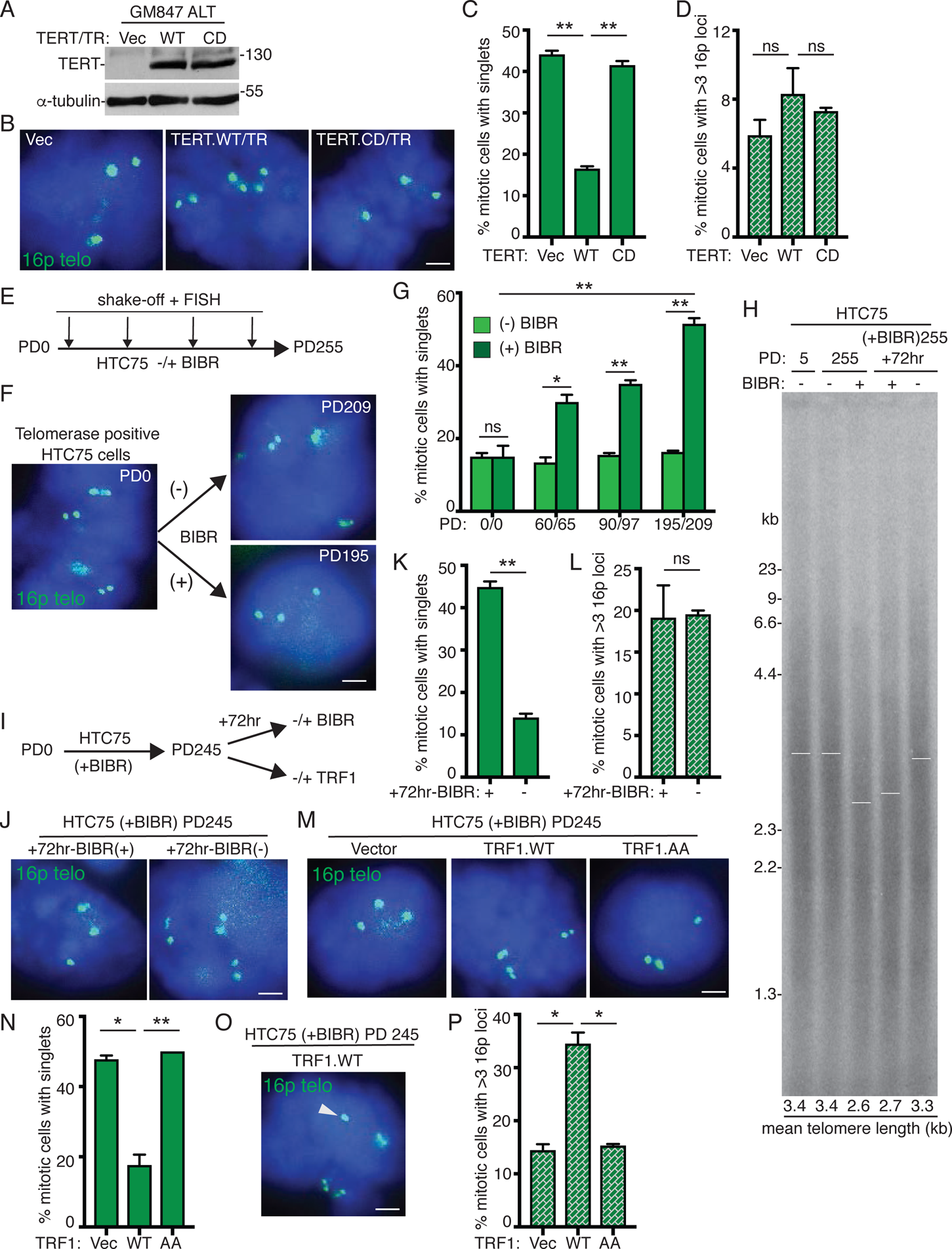
Telomerase overrides the need for persistent telomere cohesion and suppresses subtelomere recombination. **(A)** Immunoblot analysis of Vector, TERT.WT/TR, or TERT.CD/TR transfected GM847 cell extracts. **(B)** FISH analysis of Vector, TERT.WT/TR, or TERT.CD/TR transfected GM847 mitotic cells using a 16p (triploid in GM847 cells) telo probe (green). (C and D) Quantification of the frequency of mitotic cells (C) with cohered telomeres or (D) exhibiting subtelomere copying. Average of two independent experiments (n=40-44 cells each) ± SEM. **(E)** Schematic diagram of experimental setup for monitoring telomere cohesion in BIBR 1532-treated HTC75 cells. **(F)** FISH analysis of HTC75 mitotic cells at PD0 and upon long-term treatment (PD209) without (-) or (PD195) with (+) BIBR using a 16p (triploid in HTC75 cells) telo probe (green). **(G)** Quantification of the frequency of mitotic cells with cohered telomeres. Average of two independent experiments (n = 40-50 cells each) ± SEM. **(H)** Analysis of telomere restriction fragments isolated from BIBR-treated HTC75 cells, fractionated on agarose gel, denatured, and probed with a ^32^P-labeled CCCTAA probe. Cells were treated without (-) BIBR (PD5 and PD255) or with (+) BIBR (PD255). At PD255, BIBR-treated cells were split and grown with (+) or without (-) BIBR for 72 hr. White bars indicate mean telomere length. **(I)** Schematic diagram of experimental setup for resolving telomere cohesion in BIBR 1532-treated HTC75 cells at PD245. **(J)** FISH analysis of BIBR-treated HTC75 (PD245) mitotic cells that were split and grown with (+) or without (-) BIBR for 72 hr using a 16p telo probe (green). (K and L) Quantification of the frequency of mitotic BIBR-treated HTC75 (PD245) cells (K) with cohered telomeres or (L) exhibiting subtelomere copying upon continued treatment with (+) or without (-) BIBR for 72 hr. Average of two independent experiments (n=28-46 cells each) ± SEM. **(M)** FISH analysis of Vector, TRF1.WT, or TRF1.AA transfected BIBR-treated HTC75 (PD245) mitotic cells using a 16p telo probe (green). **(N)** Quantification of the frequency of mitotic cells with cohered telomeres. Average of two independent experiments (n=34-45 cells each) ± SEM. **(O)** FISH analysis of a TRF1.WT transfected BIBR-treated (PD245) HTC75 mitotic cell exhibiting subtelomere copying (arrowhead) using a 16ptelo probe (green). **(P)** Quantification of the frequency of mitotic cells exhibiting subtelomere copying. Average of three independent experiments (n=34-45 cells each) ± SEM. (B, F, J, M, O) DNA was stained with DAPI (blue). Scale bars represent 2 μm. *p ≤ 0.05, **p ≤ 0.01, ***p ≤ 0.001, (ns) not significant, Student’s unpaired t test. See also Figure S5. Source data are provided as a Source Data file.

The observation that persistent telomere cohesion can be rescued by introduction of telomerase suggests that persistent cohesion results directly from telomere shortening. To address this question we inhibited telomerase in a telomerase positive HTC75 cancer cell line (which like most other telomerase positive cells does not exhibit persistent telomere cohesion) with a small molecule inhibitor BIBR (1532) ^43, 44^. Cells were passaged long term (255 PDs) in the presence or absence of BIBR and telomere cohesion analyzed by 16p FISH (triploid in HTC75 cells) at early and late PDs (Fig. 6E). At the starting point (PD0) telomeres were resolved (Fig. 6F and 6G), as expected. Untreated BIBR cells showed resolved telomeres throughout the treatment period from PD0 to PD209 (Fig. 6F and 6G). By contrast, cells passaged in BIBR exhibited a gradual increase in persistent telomere cohesion (Fig. 6F and 6G) that accompanied telomere shortening, measured by telomere restriction fragment length analysis (Fig. 6H). These data indicate that persistent cohesion is induced by telomere shortening.

We next determined the consequences of forcing resolution in these BIBR-treated HTC75 cells at late PD (245). We took two approaches to force resolution of cohesion: reactivation of telomerase or overexpression of TRF1 (Fig. 6I). For reactivation of telomerase, cells were split and grown in the presence or absence of BIBR for an additional 72 hr. This treatment is sufficient to restore telomere length (Fig. 6H). FISH analysis shows that reactivation of telomerase leads to resolution of telomere cohesion (Fig. 6J and 6K), but not subtelomere recombination (Fig. 6L). Thus, similar to the results described above for ALT cells, telomerase forced resolution of cohesion, but at the same time suppressed subtelomere recombination between the resolved and extended telomeres. For the second approach, we introduced Vector, TRF1.WT, or TRF1.AA into BIBR-treated HTC75 cells at PD245 by transient transfection (Fig. S5E). TRF1.WT, but not Vector or TRF1.AA, forced resolution of telomere cohesion (Fig. 6M and 6N), similar to removal of BIBR. However, unlike removal of BIBR, TRF1.WT overexpression induced subtelomere recombination (Fig. 6O and 6P).

### Loss of persistent telomere cohesion and subtelomere recombination accompany senescence onset

Our data indicate that as telomeres shorten, persistent telomere cohesion driven by insufficient TRF1 (and lack of tankyrase 1 recruitment), protects cells from subtelomere recombination and premature activation of senescence. The question remains, what ultimately triggers the senescence program. As described above, when senescence was bypassed by introduction of SV40-LT, we observed a slight increase in subtelomere recombination, even without forcing resolution of telomere cohesion. We thus wondered if aged cells might ultimately succumb to subtelomere recombination, which would induce senescence. To address this question we passaged IMR90 cells until they senesced and then isolated mitotic cells by mechanical shake-off from the final passage (here defined by cessation of growth as PD51) and from two previous passages, and performed FISH with a 13q dual subtelomere/arm probe. As shown in Fig. 7A and 7B, at PD48 cells exhibited persistent telomere (but not arm) cohesion, as expected.

**Figure 7.**
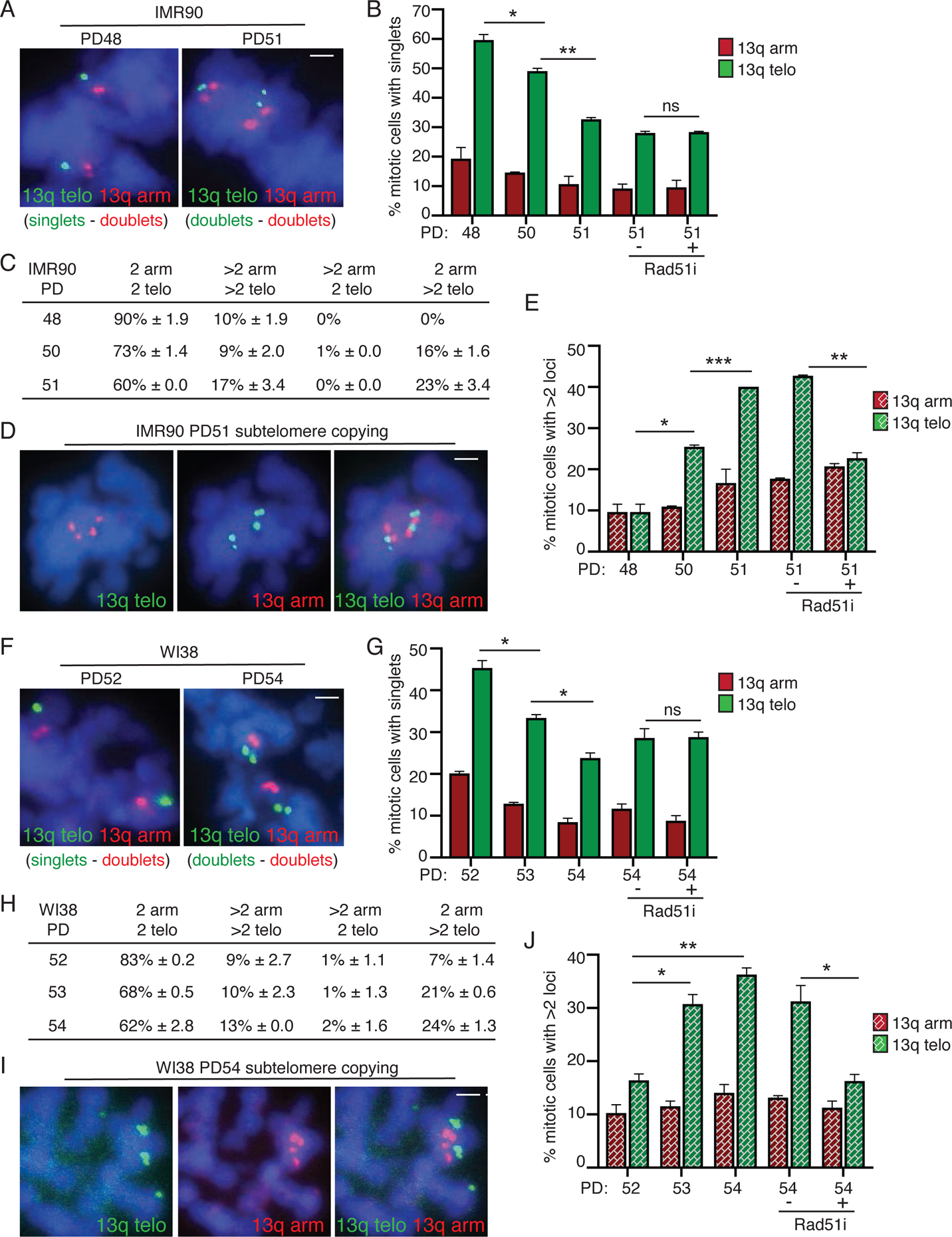
Loss of persistent telomere cohesion and subtelomere recombination accompany senescence onset. **(A)** FISH analysis of late IMR90 mitotic cells at PD48 and the final PD51 with a dual 13q arm (red) and telo (green) probe. **(B)** Quantification of the frequency of mitotic cells with cohered 13q arms (red) or telos (green). Average of two independent experiments (n=25-40 cells each) ± SEM. **(C)** Table of 13q FISH categories scored in IMR90 cells at PD48, 50 and 51. Average of two independent experiments (n=25-40 cells each) ± SEM. **(D)** FISH analysis of a final PD51 IMR90 mitotic cell exhibiting subtelomere, but not arm, copying using a dual 13q arm (red) and telo (green) probe. **(E)** Quantification of the frequency of mitotic cells exhibiting subtelomere or arm copying. Average of two independent experiments (n=25-40 cells each) ± SEM. **(F)** FISH analysis of late WI38 mitotic cells at PD52 and the final PD54 with a dual 13q arm (red) and telo (green) probe. **(G)** Quantification of the frequency of mitotic cells with cohered 13q arms (red) or telos (green). Average of two independent experiments (n=32-46 cells each) ± SEM. **(H)** Table of 13q FISH categories scored in late WI38 mitotic cells at PD 52, 53, and the final PD54. Average of two independent experiments (n=32-46 cells each) ± SEM. **(I)** FISH analysis of a final PD54 WI38 mitotic cell exhibiting subtelomere, but not arm, copying using a dual 13q arm (red) and telo (green) probe. **(J)** Quantification of the frequency of mitotic cells exhibiting subtelomere copying. Average of two independent experiments (n=32-46 cells each) ± SEM. (A, D, F, I) DNA was stained with DAPI (blue). Scale bars represent 2 μm. *p ≤ 0.05, **p ≤ 0.01, ***p ≤ 0.001, (ns) not significant, Student’s unpaired t test. Source data are provided as a Source Data file.

However, at PD50 and 51 we observed a significant reduction in persistent telomere cohesion (Fig. 7A and 7B). We next asked if this natural loss of cohesion was accompanied by subtelomere copying. We performed FISH analysis and scored four categories of 13q FISH signals: 2arm/2telo (normal ploidy); >2arm/>2telo (aneuploidy); >2arm/2telo (arm copying); and 2arm/>2telo (telomere copying) (Fig. 7C). We observed an increase in ploidy as has been described previously ^45^. In addition, we observed a significant increase in subtelomere (but not arm) copying at PD50 and 51 that was abrogated by RAD51 inhibition (Fig. 7D-7E). Similar results were obtained with WI38 cells at their final passage (defined by cessation of growth as PD54) (Fig. 7F-7J). Together these data indicate that as telomeres shorten and cells approach senescence, transient persistent telomere cohesion protects cells from subtelomere recombination and premature senescence, but ultimately telomere erosion results in loss of telomere cohesion, RAD51-dependent subtelomere recombination, and senescence activation.

## Discussion

Telomere shortening serves as a molecular clock to count the cell divisions leading to replicative senescence. Concomitant with loss of TTAGGG repeats is loss of the shelterin TTAGGG-repeat binding proteins TRF1 and TRF2. Previous studies show how loss of TRF2 could signal senescence onset ^22, 23, 26^. Here we show the impact of TRF1 loss. As telomeres shorten, the inability to recruit sufficient TRF1 and as a result tankyrase 1 leads to persistent telomere cohesion in mitosis that protects shortened telomeres from inappropriate recombination and DNA damage-induced senescence. This persistent cohesion however only affords transient protection; ultimately telomeres are unable to maintain cohesion and cells succumb to subtelomere recombination and growth arrest.

Persistent cohesion appears to be an ineluctable consequence of telomere shortening. It occurs in normal human cells as they approach senescence and their telomeres become critically short and in ALT cells, which lack telomerase and exhibit critically short telomeres at high frequency. In both cases, the observation that it can be rescued by telomerase expression indicates telomere shortening as the basis.

Conversely, our demonstration that persistent cohesion can be induced in telomerase positive cancer cells through telomerase inhibition supports the notion that persistent cohesion is inherent to short telomeres.

We showed that persistent cohesion protects cells from illegitimate subtelomere recombination. Upon forced resolution of cohesion, subtelomere recombination occurs rapidly, at high frequency, and across multiple cell types. Recombination can be induced at similar frequency in aged cells, ALT cancer cells, and telomerase-inhibited telomerase positive cancer cells. A common feature of these cells is that their telomeres have gone through many rounds of division in the absence of telomerase. Such telomeres are likely to have a subpopulation that are critically short. In the absence of telomerase, telomeres may accumulate single stranded DNA and may be eroded into the subtelomeres and thereby likely to be engaged in strand invasion and copying. Persistent cohesion, which has been shown to promote recombination between sisters in normal human cells ^46^ and in ALT cells ^31^, may benefit critically short telomeres by permitting repair, while at the same time preventing release and engagement of recombination with non-sisters. The observation that telomerase expression (when coupled to forced resolution of cohesion) abrogates subtelomere recombination, indicates short telomeres as the basis.

Although subtelomere copying in human cells was first detected in ALT, we now demonstrate that the phenomenon is not unique to ALT cells; it can occur in any cell type that lacks telomerase and has shortened telomeres. The dependence on ATR, CHK1, and RAD51 indicate that it may rely on mechanisms that facilitate long range telomere movement ^47^. The lack of dependence on RAD52 and POLD3 indicate that it is distinct from DNA damage-induced mechanisms of telomere synthesis in ALT ^36–39^. In fact, in contrast to mechanisms of telomere recombination in ALT cells, which take advantage of elevated levels of DNA damage at telomeres to promote telomere recombination for telomere extension and cell growth, subtelomere copying actually induces DNA damage and halts cell growth. Thus, persistent cohesion (in any cell type) likely serves a protective role by providing shortened (perhaps endogenously damaged) telomeres with a sister for DNA repair and by preventing damage-inducing subtelomere recombination with non-sisters.

We show that persistent cohesion protects aged cells from premature senescence, but ultimately at the final division cohesion is lost, subtelomere copying ensues, and cells senesce. Maintenance of telomere cohesion requires the shelterin subunit TIN2. Thus, it is likely that critically short telomeres lacking both TRF1 and TRF2 can no longer recruit sufficient TIN2 to maintain cohesion, thereby unleashing subtelomere recombination and DNA damage. Indeed, we showed that senescence bypass by inactivation of the DNA damage checkpoint leads to a rapid and dramatic increase in subtelomere copying. Thus, when telomere shortening is unchecked, subtelomere recombination ensues. Ultimately, cells will go through crisis and only those that develop telomere maintenance mechanisms will survive. In the case of ALT cancer cells, persistent cohesion may promote survival by providing a safe harbor for critically short telomeres

## Materials and Methods

### Cell lines

WI38 and IMR90 fibroblast cell lines (ATCC) were supplemented with 20% FBS and grown in standard conditions. For aged WI38, cells were analyzed at PD 50-55. For aged IMR90, cells were analyzed at PD 48-52. ALT cancer cell lines (GM847 and U2OS) and the telomerase-positive cancer cell line HTC75 ^11^ were supplemented with 10% FBS and grown in standard conditions. Where indicated, the following inhibitors were dissolved in DMSO and added to the culture media 16 hr prior to harvest: Rad51 inhibitor RI-1 (Selleckchem, 20 µM), Rad52 inhibitor AICAR (Selleckchem, 20 µM), ATR inhibitor VE-821 (Selleckchem, 10µM), CHK1 inhibitor SB 218078 (Tocris, 250 nM), and CHK2 inhibitor (Sigma, 10 µM).

### Generation of TRF1 mutant cell lines using CRISPR/Cas9

The tankyrase binding site (^13^RGCADG^18^) in the acidic domain of TRF1 was mutated using RNA-guided CRISPR associated nuclease Cas9. A 20 bp target sequence directed against the first exon of the human TRF1 gene (TERF1) (TRF1 guide DNA 5’ – CGCGGGGCTGTGCGGATGGT – 3’) was inserted into the guide sequence insertion site using Bbs1 site of the CRISPR plasmid pX330 comprised of Cas9 and a chimeric guide RNA ^48^. A single stranded DNA homology template was designed to change Gly18 to Pro (TRF1.G18P) and also to introduce a BamHI restriction site to screen the potential clones. HEK293T cells were transfected with the pX330 plasmid and homology template ssDNA as described previously ^49^.

Following transfection, cells were re-plated for single cell cloning, propagated and screened by a PCR strategy designed to screen for gain of a BamHI site in the target site. We isolated 3 independent homozygous TRF1.G18P clones (#1, #3 and #5) based on BamHI restriction digestion pattern of the PCR products. DNA sequencing chromatogram of the PCR products from all three clones confirmed the mutation.

### Plasmids

MycTRF1.WT ^50^ contains an N-terminal myc epitope tag followed by amino acids 2 to 439 in the pLPC retroviral vector ^51^. MycTRF1.AA ^18, 35^ was generated using site-directed mutagenesis (Agilent) of the MycTRF1.WT plasmid using the sense oligonucleotide 5’-CGGGGCTGTGCGGCTGCTAGGGATGCCGACCCT– 3’ and the antisense oligonucleotide 5’– AGGGTCGGCATCCCTAGCAGCCGCACAGCCCCG –3’. TRF1.WT and AA were PCR amplified and cloned into the NheI and SalI sites of pLSJH ^34^ to generate FlagTRF1.WT and FlagTRF1.AA lentiviral constructs. Telomerase was expressed using the pBabe hTERT-hTR plasmid ^52^ (a kind gift from Kathy Collins). The catalytically dead (CD) TERT.CD-TR containing a double point mutation D530A and V531I ^53^ was generated using site-directed mutagenesis of the TERT.WT/TR construct using the sense oligonucleotide 5’-TGTACTTTGTCAAGGTGGCTATAACGGGCGCGTACGACAC-3’ and the antisense oligonucleotide 5’-GGTGTCGTACGCGCCCGTTATAGCCACCTTGACAAAGTAC-3’.

### siRNA and plasmid transfection

For plasmids, cells were transfected with Lipofectamine 2000 (Invitrogen, Grand Island, NY) according to the manufacturer’s protocol for 20 hr. For siRNA, transfection was performed with Oligofectamine (Invitrogen) according to the manufacturer’s protocol for 72 hr. The final concentration of POLD3 (4390824-s21045, Invitrogen) or GFP duplex I (Dharmacon) siRNA was 100 nM. For TRF1 plasmid transfection of siRNA transfected cells, 20 hr prior to the end of the 72-hr siRNA incubation, cells were transfected with MycTRF1.WT using Lipofectamine 2000.

### Lentiviral Infection

Lentiviruses were produced by transfection of 293FT (Invitrogen) packaging cells with a three-plasmid system as described previously ^54, 55^. 293FT cells were seeded in a 10 cm dish at 3.34 x 10^6^ cells and 24 hr later were transfected with 2.6 µg lentiviral vector, 2.6 ug pCMVΔR.89 packaging plasmid, and 260 ng pMD.G envelope plasmid using Lipofectamine 2000 (Invitrogen) according to the manufacturer’s instructions. Lentiviral supernatants were collected at 24 and 48 hr after transfection, filtered with a 0.45 µm filter (Millipore), and frozen at −80°C. Twenty-four hr before infection, target WI38 and WI38 SV40 LT cells were seeded at a density of 1.2 x 10^6^ cells in a 10 cm dish; target U2OS cells were seeded at a density of 6 x 10^5^ in a 10 cm dish. Target cells were infected for 48 hr with lentiviral supernatants supplemented with 8 µg/ml polybrene (Sigma-Aldrich). Following a 24-hr recovery in fresh media, cells were selected in 2 µg/ml puromycin. After 24 hr of selection, cells were seeded onto cover slips. 48 hr later, cells were fixed and analyzed for damage, senescence, or Rad51+TRF2 immunofluorescence.

For growth curve analyses, target cells were infected for 48 hr. Following 24 hr of selection, cells were seeded into either 24-wells (for WI38 and WI38 SV40 LT cells) or 6-wells (for U2OS) at 10,000 cells/well in media containing 2 µg/ml puromycin. The next day (Day 0), cells were harvested and counted using a hemocytometer and used to normalize the number of cells plated. Cells were then harvested and counted every 24 hr over the next 5 days. Cell numbers were calculated as a ratio of the ‘Day 0’ counts. Three technical replicates were done for each data point.

### SV40 LT immortalization of WI38 cells

For SV40 Large T antigen infection, amphotropic retroviruses were generated by transfecting 20 µg of pBabe-neoLargeTcDNA ^56^ into Phoenix Amphotropic cells (ATCC) using calcium phosphate precipitation. Five hr following transfection, fresh media was added. 24 hr following transfection, retroviral supernatant was collected every 6 or 12 hr for 48 hr. Collected virus was filtered through a 0.45 µm filter and supplemented with 10% FBS and 4 µg/ml polybrene. Target WI38 cells were infected with retrovirus for 24 hr, followed by a 24-hr recovery in standard media. Cells were then selected in 600 µg/ml geneticin (Gibco) for long-term culturing.

### Long-term telomerase inhibition with BIBR 1532

Telomerase-positive HTC75 cells were treated with the TERT inhibitor BIBR 1532 (Selleckchem) at a final concentration of 20 µM for approximately 250 population doublings. Cells were passaged approximately every 4 days (re-plated at a 1:8 dilution), and inhibitor was freshly added at each passage.

### Preparation of cell extracts

Cells were resuspended in four volumes of TNE buffer [10 mmol/L Tris (pH 7.8), 1% Nonidet P-40, 0.15 M NaCl, 1 mmol/L EDTA, and 2.5% protease inhibitor cocktail (Sigma)] and incubated for 1 hr on ice.

Suspensions were pelleted at 8000 x g for 10 min. Equal amounts of supernatant proteins (determined by Bio-Rad protein assay) were fractionated by SDS-PAGE and analyzed by immunoblotting.

### Immunoblot analysis

Immunoblots were incubated separately with the following primary antibodies: rabbit anti-Myc sc-789 (0.1 µg/ml; Santa Cruz), mouse anti-Flag F3165 (3.8 µg/ml, Sigma), rabbit anti-TERT 375 (1:500 dilution of crude serum, raised against *Escherichia coli*-derived fusion protein containing hTERT amino acids 702-841), or mouse anti-α-tubulin ascites (1:10000-1:20000, Sigma), followed by horseradish peroxidase-conjugated donkey anti-rabbit or anti-mouse IgG (1:3000, Amersham). Bound antibody was detected with Super Signal West Pico (Thermo Scientific).

### Telomere restriction fragment analysis

Genomic DNA was isolated from DMSO- or BIBR 1532-treated HTC75 cell lines and digested with *HinfI, Alu1, MboI,* and *RsaI*. Approximately 3 µg of the digested DNA was fractionated on a 1% agarose gel. Telomeres were detected by hybridization to a ^32^P end-labeled (CCCTAA)^4^ oligonucleotide probe as previously described (Dynek and Smith, 2004).

### Fluorescence-activated cell sorter (FACS) analysis

Following trypsinization at approximately 70% confluence, cells were resuspended in PBS containing 2 mM EDTA, fixed with cold 70% (vol/vol) ethanol, treated with RNAse A (ThermoFisher, 200 µg/ml), stained with propidium iodide (50 µg/ml), and analyzed with a Becton, Dickinson FACSCalibur and CellQuest Pro software. The data were modeled using FlowJo v9 software after eliminating doublets by gating to determine relative DNA content and polyploidy. Approximately 20,000 events were analyzed for each condition.

### Chromosome-specific FISH

Cells were fixed and processed as described previously ^15^. Briefly, cells were isolated mechanically by mitotic shake-off, fixed twice in methanol:acetic acid (3:1) for 15 min, cytospun (Shandon Cytospin) at 2,000 rpm for 2 min onto slides, rehydrated in 2X SSC at 37°C for 2 min, and dehydrated in an ethanol series of 70%, 80%, and 95% for 2 min each. Cells were denatured at 75°C for 2 min and hybridized overnight at 37°C with FITC-conjugated (16ptelo or 4ptelo) or TRITC-conjugated (13qtelo) subtelomere probes, or FITC-conjugated subtelomere and TRITC-conjugated arm 13q14.3 deletion probe (13qtelo/13qarm) from Cytocell. Cells were washed in 0.4X SSC at 72°C for 2 min, and in 2X SSC with 0.05% Tween 20 at RT for 30 seconds. DNA was stained with 0.2 µg/ml DAPI. For FISH analysis of transfected WI38 and IMR90 fibroblasts, cells were treated with 50 ng/ml nocodazole (Sigma) for 16 hr prior to shake-off. Mitotic cells were scored as having telomeres cohered (singlets) if 50% or more of their loci appeared as singlets, i.e., one out of two or two out of three.

### Indirect Immunofluorescence

Cells were fixed in 2% paraformaldehyde in PBS for 10 min at RT, permeabilized in 0.5% NP-40/PBS for 10 min at RT, blocked in 1% BSA/PBS, and incubated with mouse anti-Myc clone4A6 (1.0 µg/ml; Milipore) and rabbit anti-tankyrase 1 762 (1.4 µg/ml) ^57^, mouse anti-γH2AX #05-636 (1 µg/ml; Millipore) and rabbit anti-53BP1 NB 100-304 (4 µg/ml; Novus Biologicals) or rabbit anti-TIN2 701 0.36 µg/ml ^58^, or rabbit anti-TRF1 415 0.2 µg/ml ^59^ and human anti-centromere (ACA) (1:4000) antibodies. For Rad51 and TRF2 co-immunofluorescence, cells were permeabilized in Triton X-100 buffer (0.5% Triton X-100, 20 mM Hepes-KOH at pH 7.9, 50 mM NaCl, 3 mM MgCl2, 300 mM sucrose) for 5 min at RT, fixed in 3% paraformaldehyde (in PBS, 2% sucrose) for 10 min at RT, permeabilized in Triton X-100 buffer for 10 min at RT, blocked in 1% BSA/PBS, and incubated with rabbit anti-RAD51 sc-8349 (4 µg/mL; Santa Cruz Biotechnology) and mouse anti-TRF2 IMG-124A (2.5 µg/ml, Imgenex). Primary antibodies were incubated at RT for 2 hr, followed by detection with FITC-conjugated or TRITC-conjugated donkey anti-rabbit or anti-mouse antibodies (1:100; Jackson Laboratories). DNA was stained with 0.2 µg/ml DAPI.

### Detection of Senescence-associated heterochromatin foci (SAHF)

SAHF was analyzed on coverslips processed for γH2AX and 53BP1 immunofluorescence described above. A cell was scored as SAHF-positive if its DAPI counterstain had a characteristic punctate pattern ^41^.

### Senescence associated β-galactosidase assay

For the SA-**β**-galactosidase assay ^40^, cells were fixed in 2% formaldehyde and 0.2% glutaraldehyde in PBS for 5 min, washed three times in PBS, and stained for either 4 hr (for WI38 cells) or 12 hr (for U2OS cells) at 37°C in staining solution (1 mg/mL X-gal, 150 mmol/L NaCl, 2 mmol/L MgCl2, 5 mmol/L K3Fe[CN]6, 5 mmol/L K4Fe[CN]6, and 40 mmol/L NaPi, pH 6.0).

### Image acquisition

FISH and immunofluorescence images were acquired using a microscope (Axioplan 2; Carl Zeiss, Inc.) with a Plan Apochrome 63X NA 1.4 oil immersion lens (Carl Zeiss, Inc.) and a digital camera (C4742-95; Hamamatsu Photonics). Images were acquired and processed using Openlab software (Perkin Elmer). For chromosome specific FISH, if foci fell in more than one optical plane of focus, multiple planes were merged using Openlab software. SA-β Gal staining images were imaged with simple brightfield at 20X magnification using a Zeiss AxioObserver.Z1 microscope and a Axiocam 503 camera.

### Statistical analysis

Statistical analysis was performed using Prism 8 software. Data are shown as mean ± SEM. Student unpaired *t* test was applied. P < 0.05 values were considered significant: *, P ≤ 0.05; **, P ≤ 0.01; ***, P ≤ 0.001; ****, P≤ 0.0001; ns, not significant.

### Data Availability

The authors declare that the data supporting the findings of this study are available within the paper and its supplementary information files. The Source Data underlying the following Figures: 1B, D, E, H, J; 2A-E, G, H; 3B, D, F, H, I, K, M; 4A, C-E, G, H, J: 5A, D, F-J, L, M,); 6A, C, D, G, H, K, L, N, P; and 7B, C, E, G, H, J and Supplementary Figures: S1D, F, H, I, K, L; S2A-C; S4A, C, E, F; and S5A, C-E are provided in the Source Data File. All other relevant data are available upon request.

## Acknowledgements

We thank Smith lab members and Tom Meier for critical reading of the manuscript. Research reported in this publication was supported by the National Institutes of Health under award number R01CA200751 to SS and F31CA221162 to KA.

## Supplementary Figure Legends

**Figure S1.**
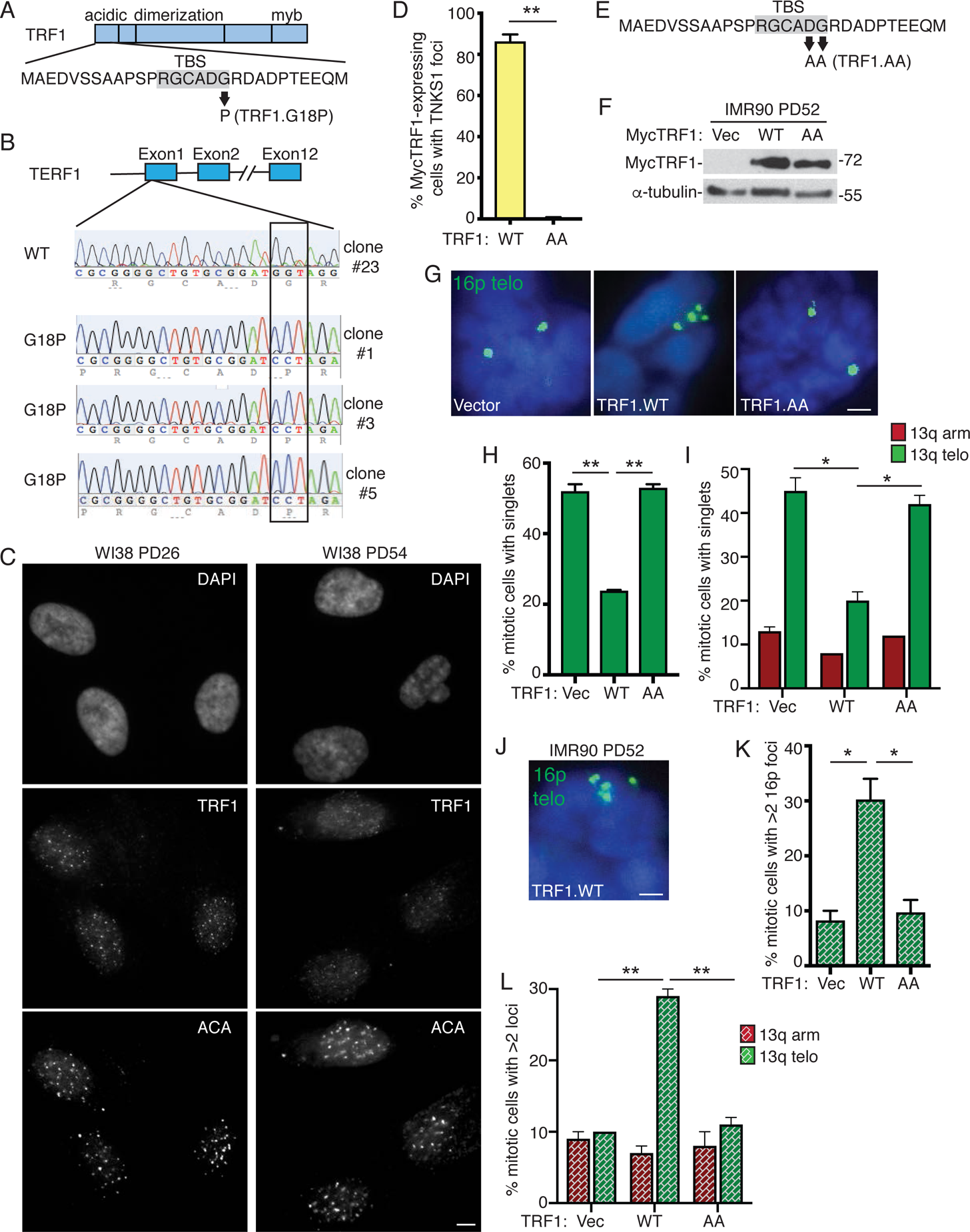
(related to. Figure 1): **(A)** Schematic diagram highlighting the G18P point mutation generated in HEK293T cells by CRISPR/Cas9 in the tankyrase-binding site (TBS) of TRF1. **(B)** Schematic diagram and corresponding sequence traces for the tankyrase-binding site of TRF1 in one wild-type HEK293T clone (#23) and three G18P mutant clones (#1, #3, #5). **(C)** Immunofluorescence analysis of early (PD26) or late (PD54) WI38 cells using anti-TRF1 or anti-centromere (ACA) antibodies. Scale bars represent 5 μm. Images were captured and reproduced under the same conditions and settings. **(D)** Quantification of the frequency of MycTRF1(WT or AA)-expressing transfected late (PD52) WI38 cells with tankyrase foci. Average of two independent experiments (n=100 cells each) ± SEM. **(E)** Schematic diagram highlighting the AA point mutation generated in MycTRF1.WT in the tankyrase-binding site (TBS). **(F)** Immunoblot analysis of Vector, TRF1.WT, or TRF1.AA transfected late (PD52) IMR90 cell extracts. **(G)** FISH analysis of Vector, TRF1.WT, or TRF1.AA transfected late (PD52) IMR90 mitotic cells using a 16p telo probe (green). **(H)** Quantification of the frequency of IMR90 mitotic cells with cohered telomeres. Average of two independent experiments (n=31-50 cells each) ± SEM. **(I)** Quantification of the frequency of mitotic cells with cohered telomeres and arms from FISH analysis of Vector, TRF1.WT, or TRF1.AA transfected late (PD50) WI38 mitotic cells using a dual 13q arm (red) and telo (green) probe. Average of two independent experiments (n=50 cells each) ± SEM. **(J)** FISH analysis of a TRF1.WT transfected late (PD52) IMR90 mitotic cell exhibiting subtelomere copying using a 16p telo probe (green). **(K)** Quantification of the frequency of mitotic cells exhibiting subtelomere copying. Average of two independent experiments (n=33-50 cells each) ± SEM. **(L)** Quantification of the frequency of mitotic cells exhibiting subtelomere copying from FISH analysis of Vector, TRF1.WT, or TRF1.AA transfected late (PD50) WI38 mitotic cells using a dual 13q arm (red) and telo (green) probe. Average of two independent experiments (n=50 cells each) ± SEM. (G and J) DNA was stained with DAPI (blue). Scale bars represent 2 μm. *p ≤ 0.05, **p ≤ 0.01, Student’s unpaired t test. Source data are provided as a Source Data file.

**Figure S2.**
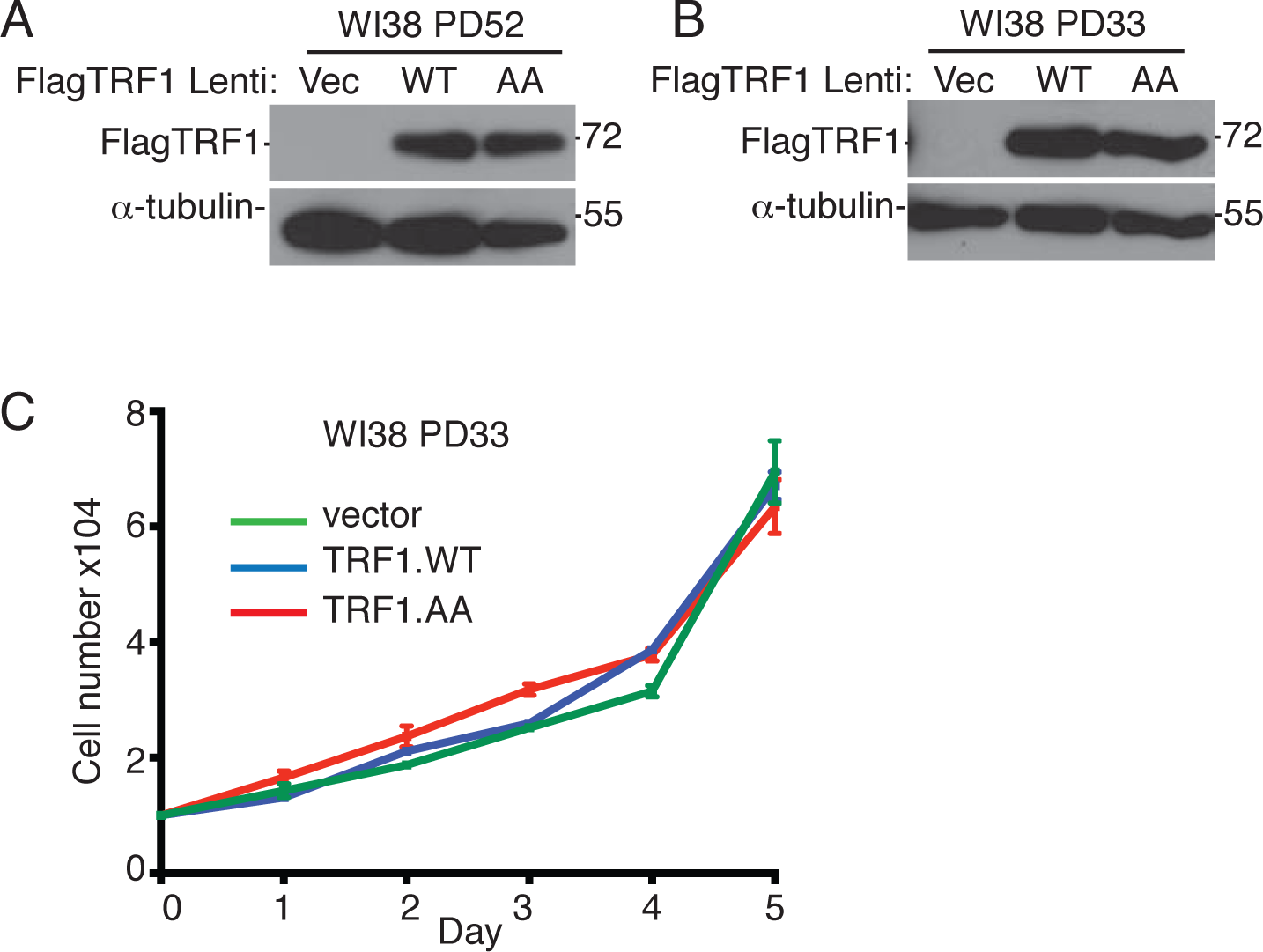
(related to Figure 3): (A and B) Immunoblot analysis of Vector, TRF1.WT, or TRF1.AA infected (A) late (PD52) or (B) early (PD33) WI38 cell extracts. (C) Growth curve analysis of Vector, TRF1.WT, or TRF1.AA infected early (PD33) WI38 cells. Average of three technical replicates ± SEM. Source data are provided as a Source Data file.

**Figure S3.**
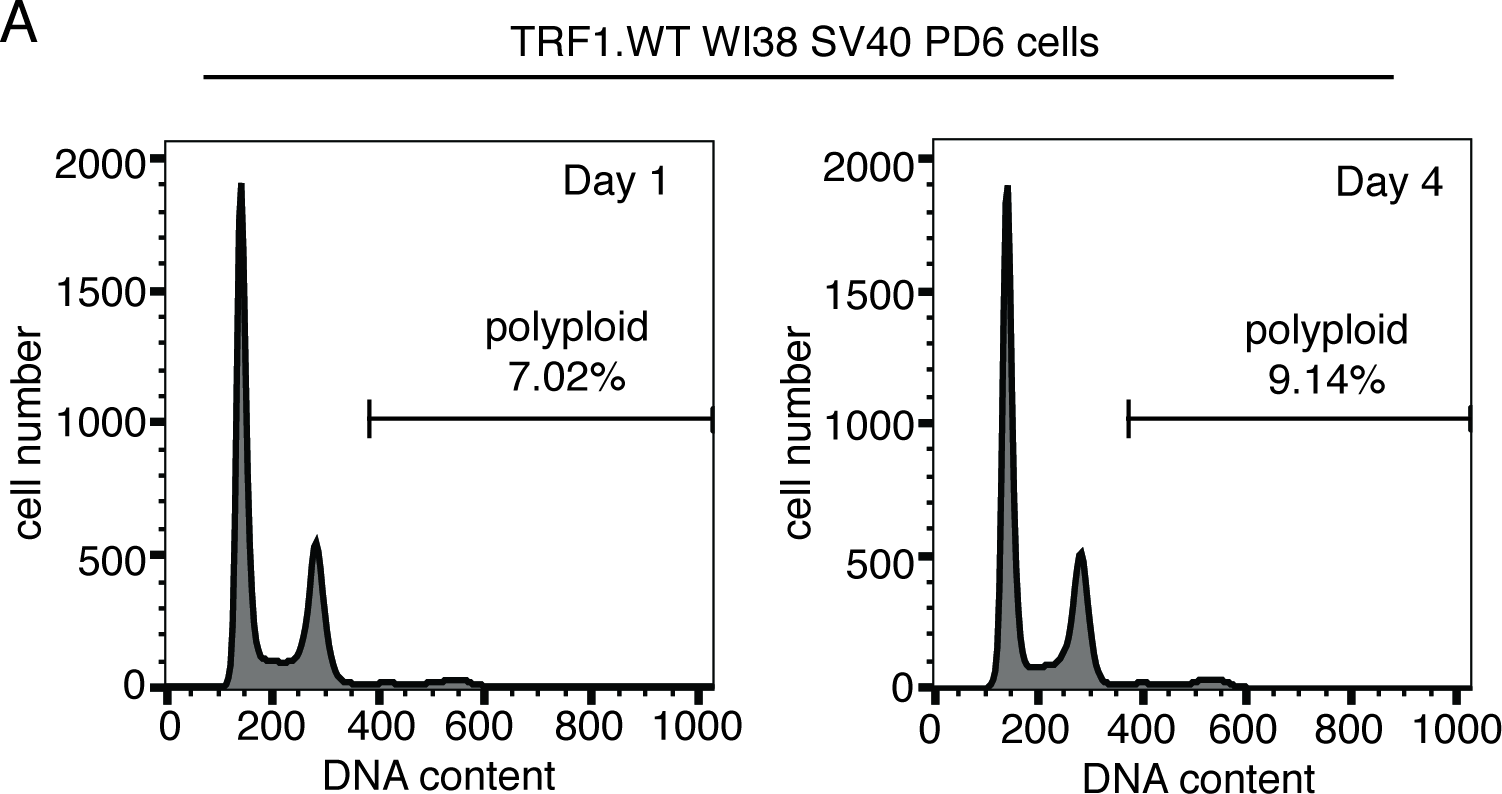
(related to Figure 4): (A) FACS cell cycle analysis of TRF1.WT infected WI38 SV40-LT (PD6) cells on Day 1 and 4 (n=approximately 20,000 events each). The percentage of events with >4N DNA content (polyploid) is shown. Source data are provided as a Source Data file.

**Figure S4.**
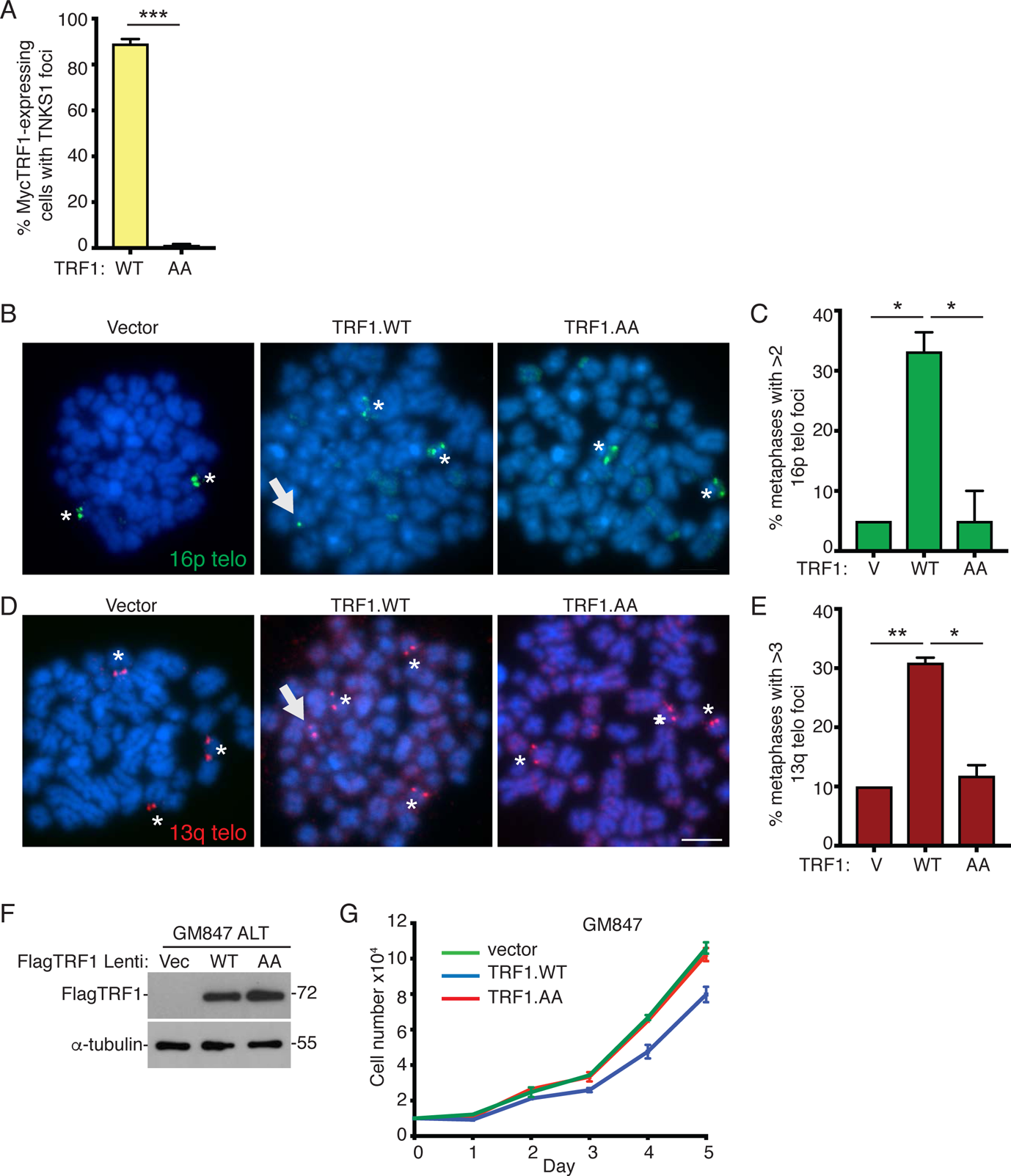
(related to Figure 5): (A) Quantification of the frequency of MycTRF1(WT or AA)-expressing transfected GM847 cells with tankyrase foci. Average of two independent experiments (n=100 cells each) ± SEM. (B and D) FISH analysis of metaphase spreads from Vector, TRF1.WT, or TRF1.AA transfected U2OS cells using (B) 16p (green) or (D) 13q (triploid in U2OS cells) (red) telo probes. Asterisks indicate usual chromosome number and arrow indicates extra subtelomere copy. (C and E) Quantification of the frequency of metaphases with an extra (C) 16p or 1(E) 3q signal. DNA was stained with DAPI (blue). Scale bar represent 5 μm. Average of two independent experiments (n=20 cells each) ± SEM. (F) Growth curve analysis of Vector, TRF1.WT, or TRF1.AA infected GM847 cells. Average of two independent experiments ± SEM. *p ≤ 0.05, **p ≤ 0.01, ***p ≤ 0.001, Student’s unpaired t test. Source data are provided as a Source Data file.

**Figure S5.**
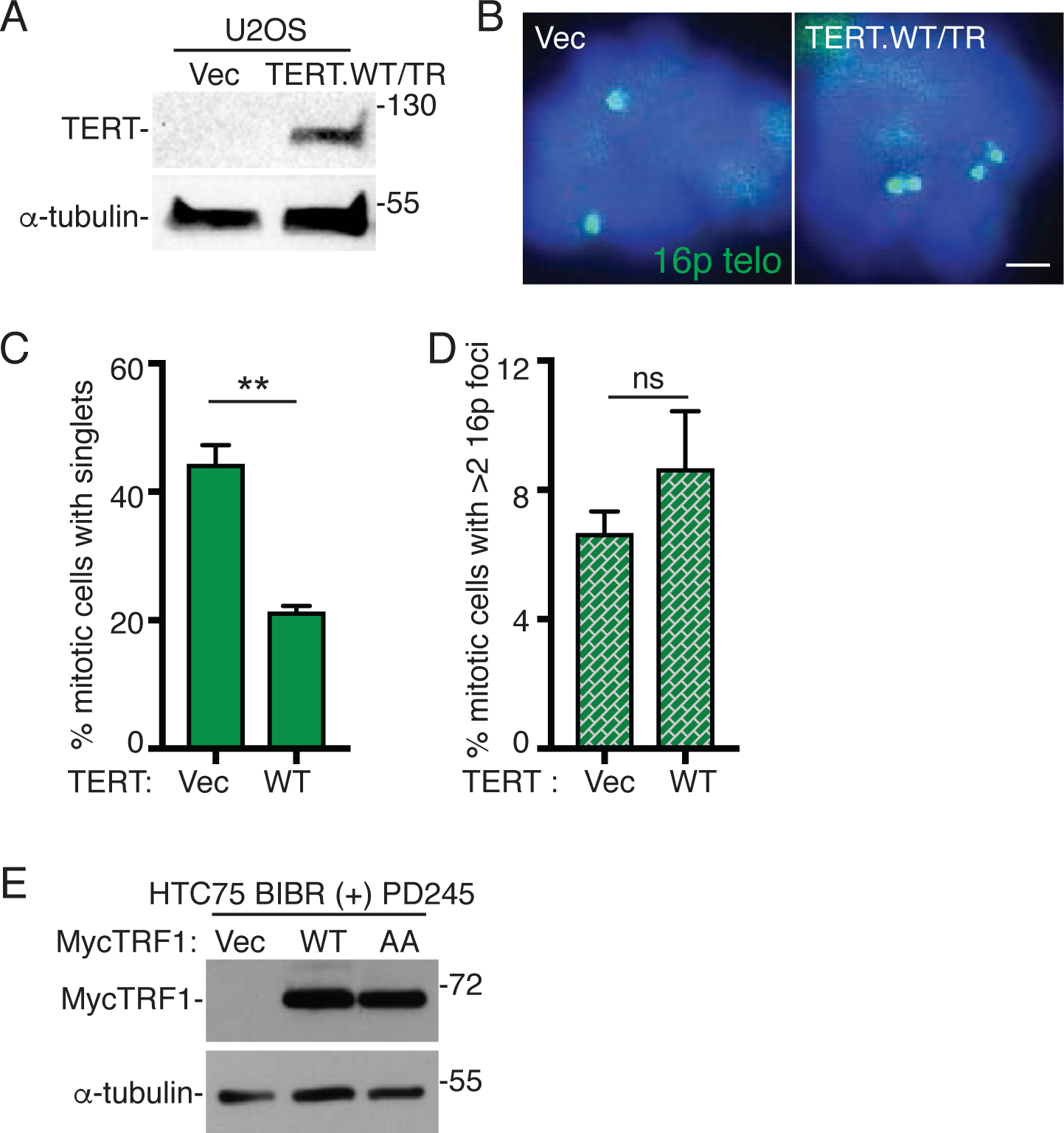
(related to Figure 6): **(A)** Immunoblot analysis of Vector and TERT.WT/TR transfected U2OS cell extracts. **(B)** FISH analysis of Vector or TERT.WT/TR transfected U2OS mitotic cells using a 16p telo probe (green). DNA was stained with DAPI (blue). Scale bar represents 2 μm. (C and D) Quantification of the frequency of mitotic cells (C) with cohered telomeres or (D) exhibiting subtelomere copying. Average of three independent experiments (n=42-45 cells each) ± SEM. (E) Immunoblot analysis of Vector, TRF1.WT, or TRF1.AA transfected HTC75 BIBR (+) PD245 cell extracts. **p ≤ 0.01, (ns) not significant, Student’s unpaired t test. Source data are provided as a Source Data file.

